# Time-resolved proteomic profile of *Amblyomma americanum* tick saliva during feeding

**DOI:** 10.1101/763029

**Authors:** Tae Kwon Kim, Lucas Tirloni, Antônio F. M. Pinto, Jolene K. Diedrich, James J. Moresco, John R. Yates, Itabajara da Silva Vaz, Albert Mulenga

**Author notes:** TKK and LT are co-first authors.

## Abstract

*Amblyomma americanum* ticks transmit more than a third of human tick-borne disease (TBD) agents in the United States. Tick saliva proteins are critical to success of ticks as vectors of TBD agents, and thus might serve as targets in tick antigen-based vaccines to prevent TBD infections. We describe a systems biology approach to identify, by LC-MS/MS, saliva proteins (tick=1182, rabbit=335) that *A. americanum* ticks likely inject into the host every 24 h during the first 8 days of feeding, and towards the end of feeding using two different sample preparation approaches (in-gel and in-solution). The in-gel approach determined molecular identification of predominant protein bands in tick saliva, and the in-solution added depth to discovery of proteins. Searching against entries in GenBank grouped tick and rabbit proteins in this study into 27 and 25 functional categories. Aside from housekeeping-like proteins, majority of tick saliva proteins belong to the tick-specific (no homology to non-tick organisms: 32%), protease inhibitors (13%), proteases (8%), glycine-rich proteins (6%) and lipocalins (4%) categories. Global secretion dynamics analysis suggests that majority (74%) of proteins in this study are associated with regulating initial tick feeding functions and transmission of pathogens as they are secreted within 24-48 h of tick attachment. Comparative analysis of the *A. americanum* tick saliva proteome to five other tick saliva proteomes identified 284 conserved tick saliva proteins: we speculate that these regulate critical tick feeding functions and might serve as tick vaccine antigens. We discuss our findings in the context of understanding *A. americanum* tick feeding physiology as a means through which we can find effective targets for a vaccine against tick feeding.

**Author Summary:** The lone star tick, *Amblyomma americanum*, is a medically important species in US that transmits 5 of the 16 reported tick-borne disease agents. Most recently, bites of this tick were associated with red meat allergies in humans. Vaccination of animals against tick feeding has been shown to be a sustainable and effective alternative to current acaricide based tick control method which has several limitations. The pre-requisite to tick vaccine development is to understand the molecular basis of tick feeding physiology. Toward this goal, this study has identified proteins that *A. americanum* ticks inject into the host at different phases of its feeding cycle. This data set has identified proteins that *A. americanum* inject into the host within 24-48 h of feeding before it starts to transmit pathogens. Of high importance, we identified 284 proteins that are present in saliva of other tick species, which we suspect regulate important role(s) in tick feeding success and might represent rich source target antigens for a tick vaccine. Overall, this study provides a foundation to understand the molecular mechanisms regulating tick feeding physiology.

## Introduction

Ticks and tick-borne diseases (TBDs) have been on the rise and have greatly impacted human and veterinary medicine. Ticks have gained the attention in public health policy with a recent publication that advocated for One Health solutions listing 17 human TBDs among sources of human health concerns (1). Moreover, the dramatic rise related to ticks and TBDs have caught the attention of United States (US) lawmakers, as shown in the 21^st^ Century Cures Act of 2016, which created the TBD Working Group. Under the Cures Act, the TBD Working Group was tasked with evaluating the impact of TBDs and required research to find solutions (https://www.hhs.gov/ash/advisory-committees/tickbornedisease/index.html). Likewise, six of the 23 human vector-borne diseases that are listed by the World Health Organization are tick borne that include Crimean-Congo haemorrhagic fever, Lyme disease, relapsing fever, rickettsial diseases (spotted fever and Q fever), tick-borne encephalitis, and tularemia (http://www.who.int/news-room/fact-sheets/detail/vector-borne-diseases). In the US, *Amblyomma americanum,* the lone star tick is among one of the tick species of medical and veterinary health significance.

*A. americanum* is a geographically expanding tick species (2) that is involved in transmission of multiple human and animal disease agents. In public health, *A. americanum* is the principal vector for *Ehrlichia chaffensis*, the causative agent of human monocytic ehrlichiosis (3), and *E. ewingii,* which also causes ehrlichiosis, referred to as human granulocytic ehrlichiosis (4–6). This tick also transmits *Francisella tularensis,* the causative agent for tularemia (7, 8), a yet to be described disease agent, suspected as *Borrelia lonestari,* which causes Lyme disease-like symptoms referred to as southern tick-associated rash illness (STARI) (9, 10) and also an *E. ruminantium-*like organism referred to as the Panola Mountain *Ehrlichia* (PME) (11). There is also evidence that *A. americanum* may transmit *Rickettsia amblyommii*, *R. rickettsia, and R. parkeri,* the causative agents to rickettsiosis to humans (12, 13). This tick has also been reported to transmit the Heartland and Bourbon viruses to humans (14, 15). Most recently, this tick has been shown to be responsible for causing an α-gal allergy or mammalian meat allergy (MMA) in humans upon tick bite (16). In veterinary health, *A. americanum* transmits *Theileria cervi* to deer (17), and *E. ewingii* to dogs (18). There are reports of mortality in deer fawns that were attributed to a combination of heavy *A. americanum* infestation and *T. cervi* infections (19). In livestock production, heavy infestations were thought to cause low productivity in cattle (20, 21). In the Southern US, *A. americanum* appears to be the most dominant tick species that bite humans, which has been reported to be responsible for 83% of human tick infestations (22).

The success of ticks as pests and vectors of TBD agents is facilitated by secreted tick salivary proteins that are injected into the host to regulate the tick’s evasion of host defense (23). There is evidence that repeatedly infested animals develop immunity against tick saliva proteins and are protected against TBD transmissions such as *Francisella tularensis* (24), *B. burgdorferi* (25–27), *Babesia* spp. (28), Thogoto virus (29), tick-borne encephalitis virus (30) and *T. parva bovis* (31). Therefore, identification of tick saliva proteins that ticks inject into the host during feeding might lead to development of tick saliva protein-based vaccines to prevent TBD infections.

The goal of this study was to utilize systems biology approach to identify proteins that *A. americanum* ticks injects every 24 h during feeding. This study builds upon our recent findings that identified *Ixodes scapularis* tick saliva proteins that are secreted every 24 h during first five days of feeding (32), partial and replete fed *Rhipicephalus microplus* (33), and replete fed adult and nymph *Haemaphysalis longicornis* (34). Others have reported proteins in saliva of replete fed *R. sanguineus* (35) and three and five day fed *Dermacentor andersoni* (36). A related study reported proteins in *Ornithodoros moubata* (soft tick) identified from saliva collected after four months from feeding (37). Most recently, the saliva proteomes of unfed *I. scapularis* and *A. americanum* exposed to different hosts have been identified (38). In this study, we report proteins that *A. americanum* ticks sequentially inject into the host every 24 h during feeding. Comparison of the *A. americanum* tick saliva proteome in this study with other saliva proteomes of other tick species allowed us to identify tick saliva proteins that are likely utilized by multiple tick species to regulate feeding, and these might represent potential antigens for anti-tick vaccine development.

## Materials and Methods

### Ethics statement

All experiments were done according to the animal use protocol approved by Texas A&M University Institutional Animal Care and Use Committee (IACUC) (AUP 2011-207 and 2011-189) that meets all federal requirements, as defined in the Animal Welfare Act (AWA), the Public Health Service Policy (PHS), and the Humane Care and Use of Laboratory Animals.

### A. americanum tick saliva collection

*A. americanum* ticks were purchased from the tick rearing facility at Oklahoma State University (Stillwater, OK, USA). Routinely, ticks were fed on rabbits as previously described (39, 40). Ticks were restricted to feed on the outer part of the ear of New Zealand rabbits with orthopedic stockinet’s glued with Kamar adhesive (Kamar Products Inc., Zionsville, IN, USA). To stimulate female *A. americanum* ticks to attach onto the host to start feeding and to be inseminated to complete the feeding process, male ticks (15 per ear) were pre-fed for three days prior to placing female ticks onto rabbit ears to feed. A total of 50 female *A. americanum* ticks (25 per ear) were placed into tick containment apparatus on each of the three rabbits and allowed to attach.

Saliva of female *A. americanum* tick was collected as previously described (32, 33). Saliva was collected from 15 ticks fed for 24, 48, 72, 96, 120, 144, 168, and 192 h, respectively, ten ticks fully fed but not detached from the host (BD) and six ticks spontaneously detached from the host (SD). Briefly, tick saliva was collected every 15 – 30 min intervals for a period of approximately 4 h at room temperature from ticks that were previously injected with 1-3 μL of 2% pilocarpine hydrochloride in phosphate buffered saline (PBS, pH 7.4) as published by our group (32, 38).

### Identification of A. americanum tick saliva proteins by LC-MS/MS

Identification of tick saliva proteins using LC-MS/MS was done in two methods: “in-gel” digestion (GeLCMS) and “in-solution” digestion (shotgun proteomics) of tick saliva protein peptides as described (32–34). For the in-gel preparation approach, saliva from a pool of 30 ticks collected from 24, 72, 120, and 168 h fed were resolved on a Novex 4-20% Tris-Glycine SDS-PAGE gradient (Thermo, Waltham, MA, USA), stained with Coomassie Brilliant Blue, visible protein bands excised and submitted for LC-MS/MS as described (33, 41). For the in-solution digestion method, ∼4.5 μg of total tick saliva proteins (in triplicate runs using ∼1.5 μg per run) per feeding time point (24, 36, 48, 72, 96, 120, 144, 168, 192, BD, and SD) were processed for LC-MS/MS as published by our group (32–34).

### Database searching of tandem mass spectra

Proteins in *A. americanum* tick saliva were identified according to the previously described pipeline (32–34). To prepare the protein database used for protein identification, we extracted the coding sequences (CDS) from *A. americanum* transcriptomes that were assembled from Illumina sequence reads (BioProject accession # PRJNA226980) (42) using an automated pipeline in Visual Basic (Microsoft, Redmond, Washington, USA) provided Dr. Jose M. Ribeiro (NIH), based on similarities to known proteins (43). Contigs from the assembled *A. americanum* transcriptome were used to identify open reading frames (ORFs) that were larger than 50 amino acids in all six frames. The identified ORFs were subjected to blastp using several amino acid sequence databases downloaded from NCBI (non-redundant [nr] Acari and refseq-invertebrate), Uniprot (nr-Acari), MEROPS database (44), the GeneOntology (GO) FASTA subset (45) and the conserved domains database (CDD) of NCBI (46) containing the COG (47), PFAM (48), and SMART motifs (49). As a false-discovery approach to identify transcripts related to hosts, we searched the ORFs against the nr-databases from NCBI for rabbit, mouse, rat, goat, sheep, cow, monkey, and humans. CDS were extracted from blastp searches that matched with 70% identity and e-value of 1e^-40^. To remove redundancies, CD-HIT (50) was used to remove sequences at 98% identity. The extracted CDS (n=110,587) were concatenated with *Oryctolagus cuniculus* from Uniprot (www.uniprot.org) (n=21,148) and reverse sequences of all entries were used to identify peptides from tandem mass spectra.

For the in-gel method, proteins were identified by searching MS/MS spectra against the protein database (described above) using the MASCOT software version 2.2 (Matrix Science, London, UK) with the following parameters: tryptic specificity, one missed cleavage and a mass tolerance of 0.2 Da in the MS mode and 0.2 Da for MS/MS ions. Carbamidomethylation of cysteine was set as a fixed modification, and methionine oxidation was set as variable modifications. Mascot peptide identifications required ion scores higher than the associated identity scores of 20 and 35 for doubly and triply charged peptides, respectively. Protein identifications were accepted if they contained at least 2 identified peptides. To be included in this analysis, all peptide sequences had to have 100% identity with assigned proteins.

For the in-solution approach, proteins were identified by first extracting the tandem mass spectra from Thermo RAW files using RawExtract 1.9.9.2 (51) and then searching against the protein database (described above) using ProLuCID in the Integrated Proteomics Pipeline Ver.5.0.1 (52). At least two peptide matches were required to be considered a protein hit. A cutoff score was established to accept a protein false discovery rate (FDR) of 1% based on the number of decoys. Additionally, a minimum sequence length of six residues per peptide was required. Results were post processed to only accept PSMs with <10ppm precursor mass error. Finally, the protein matches from each sampled time points were concatenated into one file using Identification Compare (IDcompare) program on IP2- Integrated Proteomics Pipeline Ver.5.0.1 (52).

For functional annotation, both tick and rabbit proteins were searched against the following databases: non-redundant (NR), Acari and refseq-invertebrate from NCBI, Acari from Uniprot, MEROPS database (44), the GeneOntology (GO) FASTA subset (45), and the conserved domains database of NCBI (46) containing the COG (47), PFAM (48), and SMART motifs (49). Outputs from the blast searches were used in the classifier program in Dr. Ribeiro’s visual basic program (43) to functionally categorize the identified proteins based on the best match from among all the blast screens. The functionally annotated proteins were manually validated.

### Relative abundance and graphical visualization of secretion dynamics of A. americanum tick saliva proteins

Relative abundance and secretion dynamics were determined as described (32) using normalized spectral abundance factors (NSAF) that were validated as reliable in a label-free relative quantification approach (53–55). For each functional category or individual protein, NSAF was expressed as a percent (%) of total NSAF for that time point. Percent NSAF values were normalized using Z-score statistics using the formula 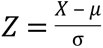, where *Z i*s the Z-score, X is the NSAF for each protein per time point, *μ* is the mean throughout time points, *σ* is the standard deviation throughout time points. Normalized percent NSAF values were used to generate heat maps using the heatmap2 function from the gplots library in R (56).

### Identification of A. americanum saliva proteins found in saliva of other tick species

*A. americanum* tick saliva proteins in this study were searched against published tick saliva proteomes of *R. microplus* (32), *I. scapularis* (33), *H. longicornis* (34), *R. sanguineus* (35), *D. andersoni* (36), and *O. moubata* (37) using local BLASTp analysis. Databases of protein sequences reported for each tick saliva proteome were extracted from NCBI or Uniprot and screened by BLASTp using the *A. americanum* saliva proteome (from this study) as the query. Protein matches ≥70% identity was reported.

## Results and Discussion

### Protein profile and abundance changes every 24 h during A. americanum tick feeding

Previous studies have demonstrated that the protein profile and abundance in salivary glands of female A. americanum is dynamic and changes during the course of tick feeding (57). However, a limitation to the previous study was that it did not inform which salivary gland proteins were secreted during feeding. To attempt at capturing changes in tick saliva protein profiles, we successfully used pilocarpine to induce and collect saliva from A. americanum ticks every 24 h during the first eight days of tick feeding as we all as from ticks that had engorged but had not detached, and replete fed ticks as described (32, 58). In early feeding stages (24-72 h), A. americanum tick saliva was observed as a white flake that accumulated on the mouthparts over time and was collected every 15 - 30 min for 4 h by washing the mouthparts with sterile phosphate buffered saline. Tick saliva was more evident after 72 h of feeding, observed as droplets of liquid forming at the mouthparts. Proteins in tick saliva were identified by LC-MS/MS sequencing in two approaches: “in-gel” digestion (GeLCMS) and “in-solution” digestion (shotgun proteomics) of tick saliva protein peptides.

For the “in-gel” digestion approach, saliva that was collected from 24, 72, 120, and 168 h fed A. americamnum ticks was electrophoresed on a 4-20% SDS-PAGE and Coomassie blue staining. Subsequently visible protein bands (n=157) (Fig. 1) were individually excised, processed for in-gel trypsinization and LC-MS/MS analysis. The peptide MS/MS spectra were searched using MASCOT software version 2.2 (Matrix Science, London, UK) against a combined protein database (tick, rabbit, and human contaminants [i.e keratin]) that was translated from coding domains (n=110,587) that were assembled from Bioproject # PRJNA226980 (42). This analysis identified a total of 76 proteins (294 peptides) in tick saliva of which 55 (229 peptides) and 21 (64 peptides) belonged to tick and rabbit, respectively (Supplemental table 1). Of the total 55 tick saliva proteins 23, 16, 41, and 19 were identified in saliva of 24, 72, 120 and 168 h fed, respectively (Tables 1A-C). Likewise, we identified 1, 19, 8, and 4 rabbit proteins in 24, 72, 120 and 168 h fed tick saliva, respectively (Tables 1A-C).

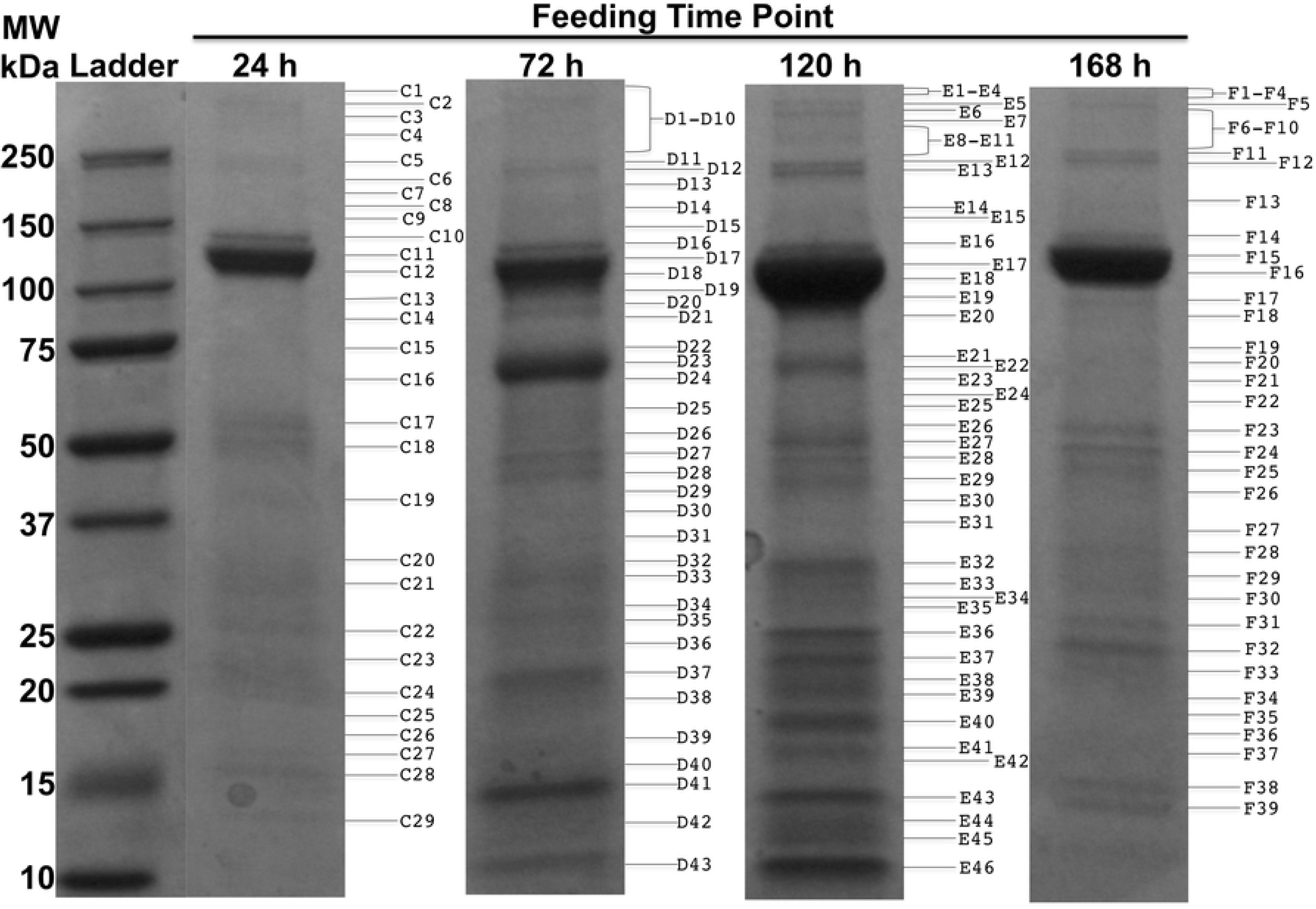

**Table 1A.**
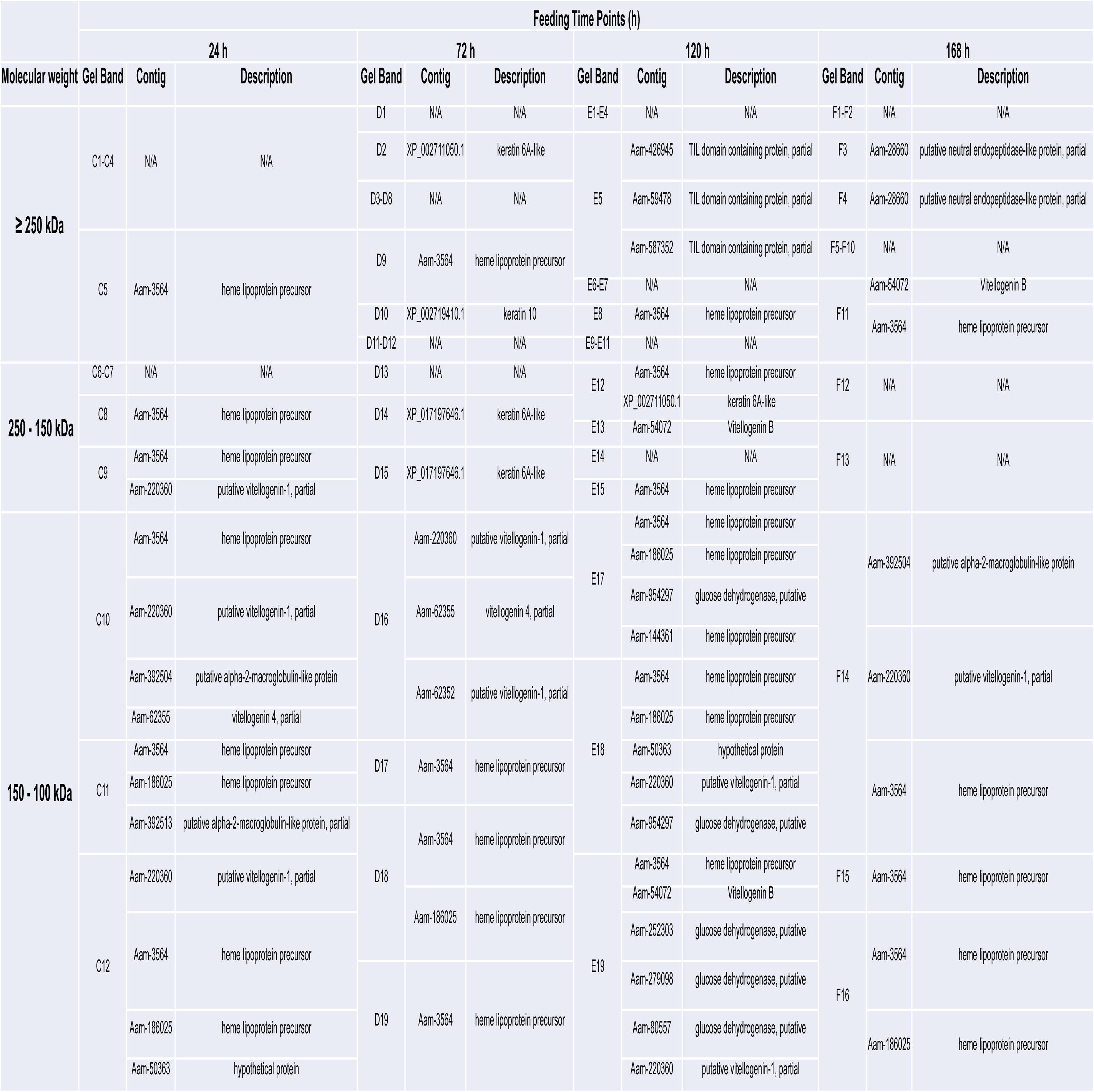
*Amblyomma america*num saliva proteins (250-100 kDa) identified from in gel digestion and LC-MS/MS during feeding

**Table 1B.**
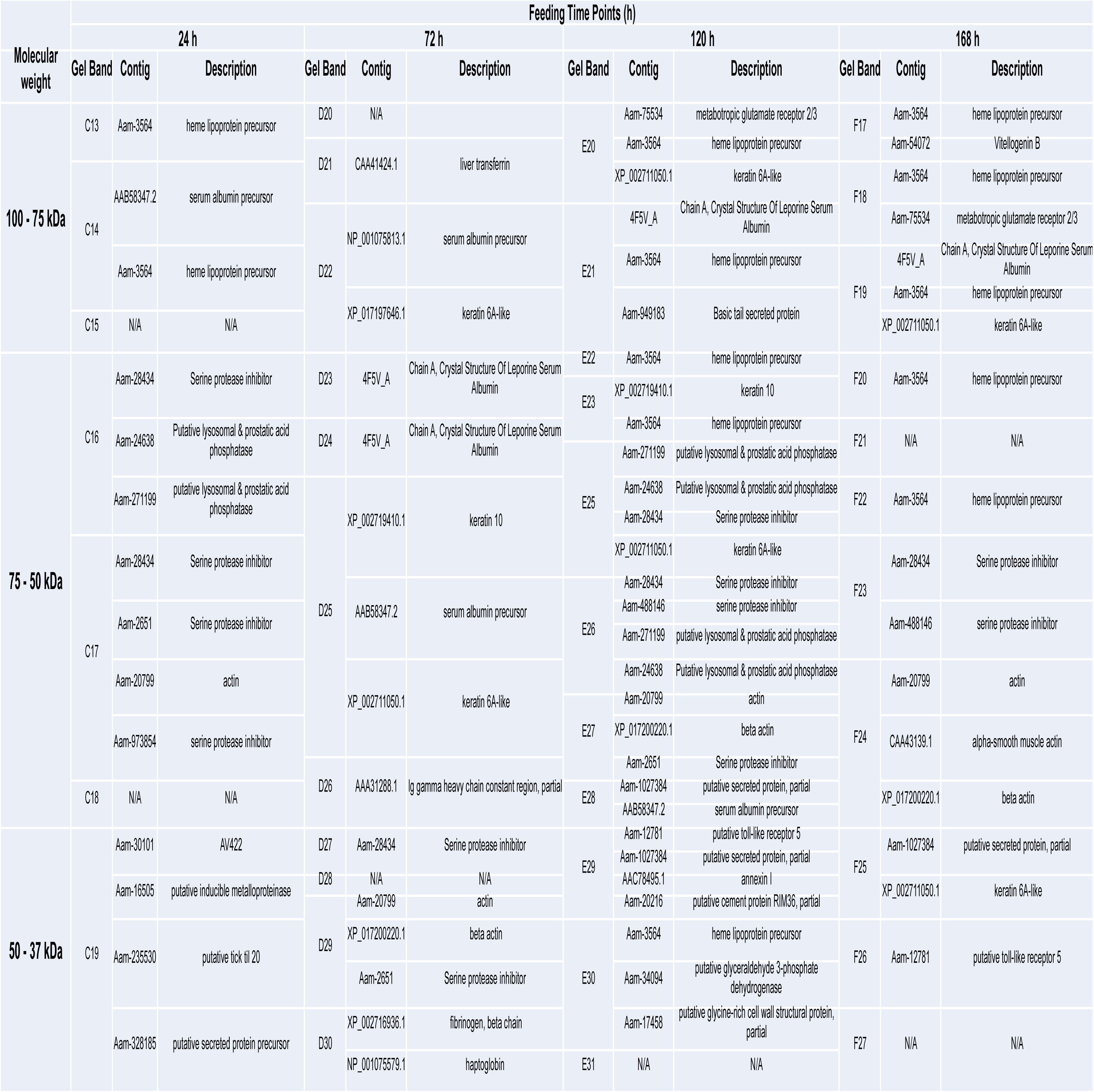
*Amblyomma america*num saliva proteins (100-37 kDa) identified from in gel digestion and LC-MS/MS during feeding

**Table 1C.**
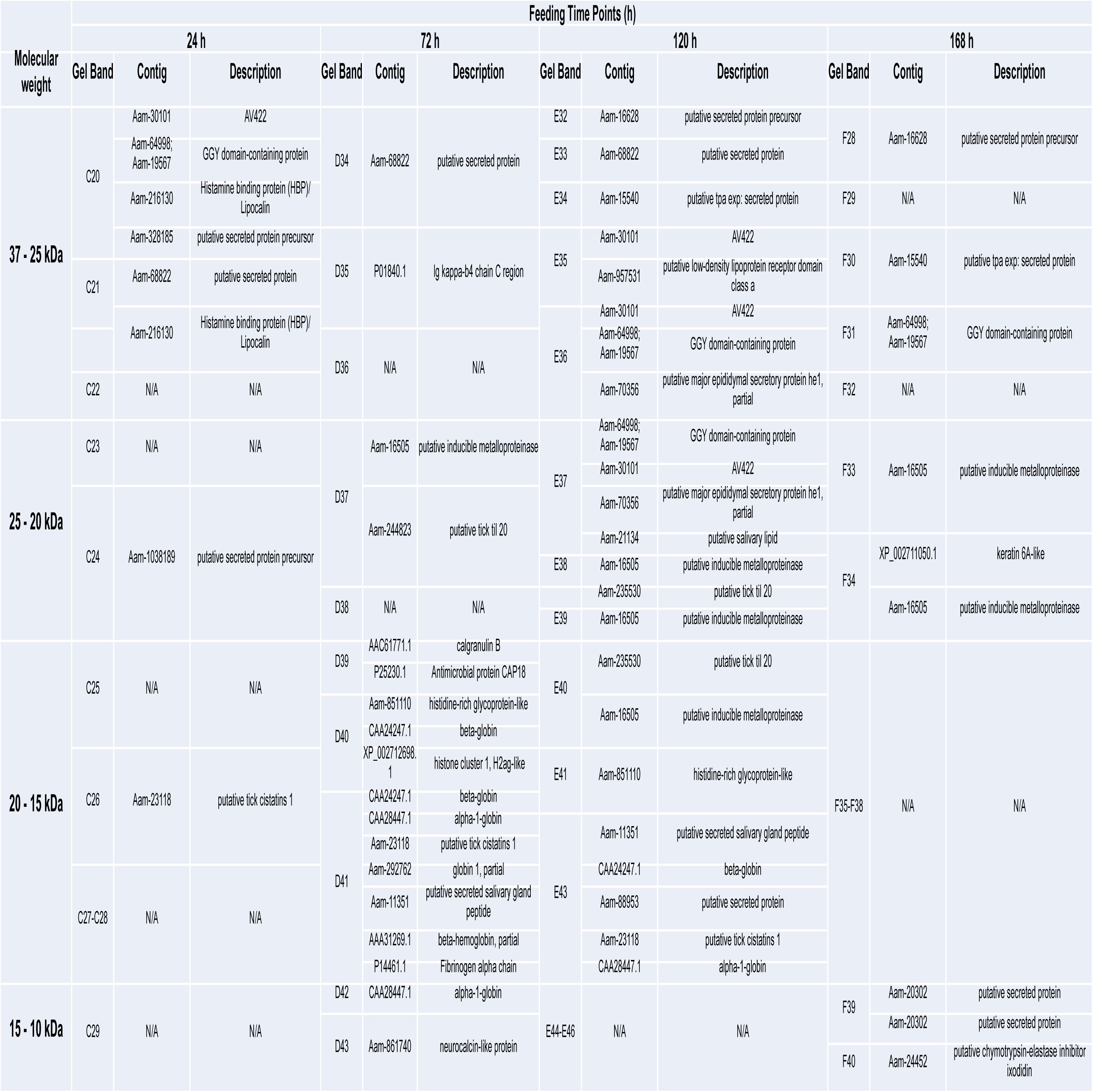
*Amblyomma america*num saliva proteins (37-10 kDa) identified from in gel digestion and LC-MS/MS during feeding

For in the “in-solution” digestion approach, saliva collected from ticks that had fed for 24, 48, 72, 96, 120, 144, 168, and 192 h as well as ticks that were apparently engorged but were not detached from the host (BD) and replete fed (SD) was subjected to LC-MS/MS analysis. Peptide mass spectra were searched against the combined database (described above) using the ProLuCID search engine (52). This analysis identified a total of 1612 proteins of which 1182, 335, 30, and 65 were considered as respective tick, rabbit, and contaminants or reversed proteins (Supplemental table 2). We respectively identified 450, 540, 419, 441, 332, 529, 478, 536, 312, and 325 tick proteins in the 10 different saliva samples. Similarly, we respectively identified 127, 130, 115, 147, 112, 140, 199, 198, 78, and 282 as rabbit proteins (Supplemental table 2). The identification of 1182 tick and 335 rabbit unique proteins in tick saliva demonstrates the complexity of tick and host interactions.

### Tick and rabbit proteins in A. americanum tick saliva are annotated in multiple functional categories

For putative functional annotation, identified proteins were searched against entries in public databases, NCBI, Uniprot, and MEROPS. This analysis categorized the 55 tick and 21 rabbit proteins that were identified in the “in-gel” digestion method into 12 and 9 functional protein categories, respectively (Tables 1A, 1B and 1C). Proteins that were identified in the “in-solution” digestion approach (1182 tick and 335 rabbit proteins) were categorized into a respective 27 (Tables 2A and 2B) and 25 (Tables 3A and 3B) functional categories. It is interesting but not surprising to note that all proteins that were identified in the “in-gel” digestion approach were among those that were identified in the “in-solution” digestion approach. We are aware of the fact that, utilizing both methods might be perceived as redundant, however the advantage of the “in-gel” digestion method was that, we determined the molecular identities of the predominant protein bands on Coomassie blue-stained A. americanum tick saliva SDS-PAGE (Fig. 1, Tables 1A-C). For instance, the predominant protein band between 100-150 kDa (Fig. 1) consists of heme-binding storage proteins called hemelipoproteins and vitellogenins.

Along with findings from electrophoretic profile, the “in-solution” approach confirmed the complexity of tick saliva. With redundancy removed at 98% amino acid identity levels, the majority of the identified proteins are tick specific proteins (did not match to proteins in non-tick organisms) of unknown function (32%), followed by protease inhibitors (PI) (13%), proteases (8%), and glycine-rich proteins (6%). Notable protein categories that were ≤ 5% include cytoskeletal, lipocalin, antioxidant/detoxification, extracellular matrix, immune related, heme/iron-binding, mucins, evasins, antimicrobials, and ixodegrins (Tables 2A and 2B, Supplemental table 2). For rabbit proteins, the majority are categorized as cytoskeletal (19%), followed by keratin (13%), nuclear regulation (8%), immunity-related (8%), globin/RBC degradation (6%), and protein categories that were ≤ 5% include antimicrobials, heme/iron-binding, protease inhibitors, proteases, extracellular matrix, antioxidant/detoxification, fibrinogen and lipocalin (Tables 3A and 3B, Supplemental table 2).

**Table 2A.** Numbers and cumulative relative abundance of tick protein classes in *Amblyomma americanum* saliva during 24-120 h of feeding

| Classification | Feeding Time Point |  |  |  |  |  |  |  |  |  |
| --- | --- | --- | --- | --- | --- | --- | --- | --- | --- | --- |
|  | 24 h |  | 48 h |  | 72 h |  | 96 h |  | 120 h |  |
|  | Protein Count | NSAF (%) | Protein Count | NSAF (%) | Protein Count | NSAF (%) | Protein Count | NSAF (%) | Protein Count | NSAF (%) |
| antimicrobial | 6 | 4.20 | 6 | 3.30 | 5 | 6.01 | 5 | 6.15 | 5 | 6.73 |
| cytoskeletal | 42 | 4.77 | 16 | 1.32 | 25 | 2.13 | 14 | 1.86 | 6 | 0.71 |
| detoxification | 19 | 2.16 | 15 | 0.71 | 12 | 0.72 | 15 | 0.89 | 3 | 0.18 |
| evasin | 0 | 0.00 | 6 | 0.74 | 2 | 0.24 | 6 | 0.68 | 7 | 1.67 |
| extracellular matrix | 20 | 1.81 | 11 | 1.20 | 17 | 1.48 | 11 | 0.64 | 6 | 0.37 |
| glycine rich | 23 | 2.41 | 43 | 5.81 | 44 | 9.82 | 41 | 4.86 | 27 | 6.83 |
| heme/iron binding | 16 | 20.20 | 16 | 7.48 | 15 | 15.27 | 16 | 12.35 | 9 | 5.81 |
| immune-related | 11 | 2.96 | 9 | 2.28 | 8 | 4.12 | 9 | 4.47 | 7 | 3.54 |
| ixodegrin | 1 | 0.46 | 5 | 0.82 | 1 | 0.84 | 2 | 0.47 | 2 | 0.62 |
| lipocalin | 8 | 0.74 | 17 | 2.90 | 2 | 0.55 | 6 | 1.01 | 3 | 4.39 |
| metabolism, amino acid | 3 | 0.08 | 0 | 0.00 | 1 | 0.02 | 0 | 0.00 | 0 | 0.00 |
| metabolism, carbohydrate | 11 | 0.57 | 7 | 0.22 | 8 | 0.36 | 9 | 0.77 | 3 | 0.06 |
| metabolism, energy | 18 | 1.22 | 17 | 1.54 | 13 | 1.48 | 10 | 0.94 | 2 | 0.04 |
| metabolism, lipid | 14 | 1.07 | 19 | 1.14 | 18 | 1.19 | 14 | 1.33 | 8 | 0.57 |
| metabolism, nucleic acid | 7 | 2.48 | 18 | 1.99 | 9 | 0.97 | 9 | 2.19 | 13 | 1.22 |
| mucin | 3 | 0.21 | 5 | 0.53 | 4 | 0.54 | 6 | 0.96 | 4 | 0.27 |
| nuclear regulation | 7 | 0.39 | 7 | 0.45 | 8 | 1.04 | 7 | 1.24 | 7 | 1.00 |
| protease | 17 | 0.72 | 25 | 1.07 | 20 | 0.97 | 25 | 1.46 | 27 | 4.08 |
| protease inhibitor | 55 | 22.28 | 76 | 21.14 | 52 | 11.10 | 64 | 15.33 | 66 | 20.32 |
| proteasome machinery | 8 | 1.57 | 6 | 0.50 | 6 | 1.29 | 6 | 1.50 | 6 | 1.18 |
| protein modification | 16 | 0.59 | 6 | 0.14 | 16 | 0.84 | 7 | 0.22 | 5 | 0.13 |
| protein synthesis | 6 | 0.16 | 2 | 0.05 | 4 | 0.14 | 2 | 0.06 | 0 | 0.00 |
| signal transduction | 14 | 3.57 | 11 | 4.74 | 3 | 0.39 | 7 | 2.01 | 8 | 1.80 |
| tick specific proteins | 110 | 23.99 | 188 | 39.44 | 118 | 37.88 | 140 | 37.87 | 105 | 38.28 |
| transcription machinery | 4 | 0.24 | 1 | 0.03 | 3 | 0.18 | 3 | 0.11 | 0 | 0.00 |
| transporter/ receptor | 10 | 1.12 | 6 | 0.42 | 4 | 0.37 | 7 | 0.62 | 3 | 0.19 |
| transposon element | 1 | 0.02 | 2 | 0.03 | 1 | 0.05 | 0 | 0.00 | 0 | 0.00 |
| Total | 450 | 100.00 | 540 | 100.00 | 419 | 100.00 | 441 | 100.00 | 332 | 100.00 |

**Table 2B.** Numbers and cumulative relative abundance of tick protein classes in *Amblyomma americanum* saliva during 144 to completion of feeding

| Classification | Feeding Time Point |  |  |  |  |  |  |  |  |  |
| --- | --- | --- | --- | --- | --- | --- | --- | --- | --- | --- |
|  | 144 h |  | 168 h |  | 192 h |  | BD |  | SD |  |
|  | Protein Count | NSAF (%) | Protein Count | NSAF (%) | Protein Count | NSAF (%) | Protein Count | NSAF (%) | Protein Count | NSAF (%) |
| antimicrobial | 4 | 4.31 | 4 | 2.87 | 3 | 2.98 | 3 | 5.04 | 3 | 1.49 |
| cytoskeletal | 38 | 3.43 | 11 | 0.98 | 16 | 1.23 | 3 | 0.39 | 14 | 1.42 |
| detoxification | 21 | 1.71 | 11 | 0.67 | 18 | 1.13 | 7 | 0.79 | 12 | 0.55 |
| evasin | 5 | 1.54 | 7 | 3.40 | 6 | 3.39 | 1 | 0.83 | 5 | 2.86 |
| extracellular matrix | 18 | 1.61 | 7 | 0.38 | 19 | 0.85 | 14 | 1.75 | 0 | 0.00 |
| glycine rich | 14 | 1.03 | 19 | 2.00 | 31 | 2.79 | 9 | 0.47 | 8 | 0.56 |
| heme/iron binding | 17 | 22.96 | 14 | 7.31 | 16 | 12.38 | 17 | 26.07 | 10 | 7.23 |
| immune-related | 14 | 3.76 | 9 | 2.37 | 14 | 3.09 | 12 | 4.65 | 7 | 1.66 |
| ixodegrin | 1 | 0.58 | 1 | 0.20 | 1 | 0.62 | 2 | 0.78 | 0 | 0.00 |
| lipocalin | 8 | 1.11 | 24 | 6.57 | 20 | 5.69 | 20 | 4.07 | 26 | 14.76 |
| metabolism, amino acid | 4 | 0.07 | 2 | 0.07 | 1 | 0.04 | 0 | 0.00 | 0 | 0.00 |
| metabolism, carbohydrate | 22 | 1.30 | 4 | 0.12 | 14 | 0.90 | 14 | 2.56 | 5 | 0.17 |
| metabolism, energy | 13 | 0.81 | 9 | 0.34 | 10 | 0.50 | 2 | 0.30 | 6 | 0.15 |
| metabolism, lipid | 16 | 0.95 | 5 | 0.39 | 16 | 1.01 | 12 | 1.29 | 3 | 0.39 |
| metabolism, nucleic acid | 14 | 1.80 | 10 | 0.93 | 16 | 1.72 | 3 | 1.11 | 8 | 1.21 |
| mucin | 7 | 0.33 | 3 | 0.13 | 9 | 0.33 | 3 | 0.56 | 0 | 0.00 |
| nuclear regulation | 10 | 0.74 | 8 | 1.49 | 8 | 1.02 | 1 | 0.26 | 11 | 2.15 |
| protease | 40 | 3.37 | 38 | 4.57 | 43 | 4.21 | 36 | 2.93 | 36 | 7.70 |
| protease inhibitor | 83 | 17.41 | 81 | 15.82 | 78 | 14.79 | 59 | 19.17 | 39 | 17.08 |
| proteasome machinery | 7 | 0.87 | 6 | 1.09 | 6 | 0.87 | 4 | 0.11 | 6 | 1.68 |
| protein modification | 18 | 0.58 | 10 | 0.26 | 13 | 0.38 | 0 | 0.00 | 11 | 0.44 |
| protein synthesis | 2 | 0.03 | 2 | 0.06 | 5 | 0.15 | 0 | 0.00 | 2 | 0.08 |
| signal transduction | 14 | 3.11 | 10 | 1.49 | 9 | 2.43 | 7 | 1.89 | 5 | 0.53 |
| tick specific proteins | 127 | 25.15 | 175 | 46.18 | 157 | 36.86 | 77 | 23.84 | 103 | 37.75 |
| transcription machinery | 2 | 0.04 | 1 | 0.02 | 1 | 0.01 | 0 | 0.00 | 1 | 0.02 |
| transporter/ receptor | 10 | 1.38 | 6 | 0.26 | 6 | 0.62 | 6 | 1.14 | 3 | 0.07 |
| transposon element | 0 | 0.00 | 1 | 0.04 | 0 | 0.00 | 0 | 0.00 | 1 | 0.02 |
| Total | 529 | 100.00 | 478 | 100.00 | 536 | 100.00 | 312 | 100.00 | 325 | 100.00 |

**Table 3A.** Numbers and cumulative relative abundance of rabbit protein classes in *Amblyomma americanum* saliva during 24-120 h of feeding

|  | Feeding Time Point |  |  |  |  |  |  |  |  |  |
| --- | --- | --- | --- | --- | --- | --- | --- | --- | --- | --- |
|  | 24 h |  | 48 h |  | 72 h |  | 96 h |  | 120 h |  |
| Classification | Protein Count | NSAF (%) | Protein Count | NSAF (%) | Protein Count | NSAF (%) | Protein Count | NSAF (%) | Protein Count | NSAF (%) |
| Antimicrobial | 3 | 0.00134357 | 4 | 0.00192864 | 3 | 0.00171397 | 5 | 0.0086771 | 6 | 0.00763598 |
| Antioxidant | 2 | 0.00042155 | 1 | 7.576E-05 | 1 | 0.00035322 | 0 | 0 | 0 | 0 |
| Cytoskeletal | 19 | 0.01798265 | 14 | 0.01629188 | 11 | 0.00953933 | 12 | 0.01698913 | 7 | 0.01026072 |
| Extracellular matrix | 3 | 0.00049492 | 1 | 0.000267 | 1 | 0.0000291 | 2 | 0.0001466 | 2 | 0.0003605 |
| Fibrinogen | 3 | 0.00022576 | 4 | 0.00017475 | 0 | 0 | 1 | 6.7612E-05 | 2 | 0.00022745 |
| Globin/ RBC | 20 | 0.08991802 | 18 | 0.02390767 | 12 | 0.01107268 | 15 | 0.02658361 | 15 | 0.02722383 |
| Heme/Iron binding | 5 | 0.01948816 | 7 | 0.01314713 | 2 | 0.00443612 | 7 | 0.00916951 | 4 | 0.00596773 |
| Immunity related | 9 | 0.00268497 | 7 | 0.0023492 | 5 | 0.00211006 | 8 | 0.00712898 | 15 | 0.01097912 |
| Keratin | 17 | 0.00337019 | 32 | 0.01031924 | 35 | 0.02503904 | 41 | 0.04692297 | 27 | 0.03080678 |
| Lipocalin | 0 | 0 | 1 | 0.00051272 | 1 | 0.00052663 | 1 | 0.00283525 | 1 | 0.00167895 |
| Metabolism, amino acid | 0 | 0 | 0 | 0 | 0 | 0 | 0 | 0 | 0 | 0 |
| Metabolism, carbohydrates | 5 | 0.00133936 | 4 | 0.00037672 | 5 | 0.00064097 | 5 | 0.00074812 | 3 | 0.00033307 |
| Metabolism, energy | 1 | 2.8207E-05 | 0 | 0 | 0 | 0 | 3 | 0.00071594 | 1 | 0.00018482 |
| Metabolism, lipid | 0 | 0 | 0 | 0 | 0 | 0 | 2 | 0.00093604 | 2 | 0.00138574 |
| Metabolism, nucleic acids | 0 | 0 | 0 | 0 | 0 | 0 | 0 | 0 | 0 | 0 |
| Nuclear regulation | 8 | 0.00325694 | 16 | 0.00871704 | 15 | 0.01681692 | 18 | 0.01938734 | 15 | 0.01609353 |
| Protease | 0 | 0 | 1 | 0.00013596 | 0 | 0 | 2 | 0.00146001 | 1 | 4.2416E-05 |
| Protease Inhibitors | 8 | 0.0025413 | 5 | 0.00116042 | 1 | 0.0001394 | 5 | 0.00178596 | 1 | 0.00071414 |
| Protein export | 0 | 0 | 2 | 0.00017604 | 2 | 0.00115523 | 3 | 0.00209215 | 3 | 0.00247782 |
| Protein modification | 7 | 0.00102993 | 3 | 0.0003923 | 7 | 0.00221869 | 6 | 0.00095378 | 4 | 0.00051132 |
| Protein synthesis | 4 | 0.00135166 | 2 | 0.0001756 | 4 | 0.00122153 | 4 | 0.0006344 | 0 | 0 |
| Proteasome machinery | 2 | 0.00600273 | 2 | 0.0021349 | 2 | 0.00578655 | 2 | 0.0056957 | 2 | 0.00490593 |
| Signal transduction | 9 | 0.00264391 | 1 | 0.00047693 | 7 | 0.00264375 | 1 | 0.00089234 | 1 | 0.00035228 |
| Transporter/ Receptor | 2 | 0.00013431 | 5 | 0.00049163 | 1 | 3.5648E-05 | 4 | 0.00061835 | 0 | 0 |
| Transcription machinery | 0 | 0 | 0 | 0 | 0 | 0 | 0 | 0 | 0 | 0 |
| Total | 127 | 0.15425811 | 130 | 0.08321153 | 115 | 0.08547884 | 147 | 0.15444091 | 112 | 0.12214214 |

**Table 3B.** Numbers and cumulative relative abundance of rabbit protein classes in *Amblyomma americanum* saliva during 144 to completion of feeding

|  | Feeding Time Point |  |  |  |  |  |  |  |  |  |
| --- | --- | --- | --- | --- | --- | --- | --- | --- | --- | --- |
|  | 144 h |  | 168 h |  | 192 h |  | BD |  | SD |  |
| Classification | Protein Count | NSAF (%) | Protein Count | NSAF (%) | Protein Count | NSAF (%) | Protein Count | NSAF (%) | Protein Count | NSAF (%) |
| Antimicrobial | 8 | 0.007565 | 9 | 0.018337 | 9 | 0.015212 | 6 | 0.004394 | 8 | 0.015088 |
| Antioxidant | 3 | 0.000677 | 2 | 0.00071 | 3 | 0.000384 | 0 | 0 | 4 | 0.000927 |
| Cytoskeletal | 20 | 0.019936 | 29 | 0.014844 | 26 | 0.019163 | 6 | 0.011922 | 55 | 0.031601 |
| Extracellular matrix | 1 | 3.63E-05 | 0 | 0 | 1 | 2.58E-05 | 1 | 8.41E-05 | 3 | 0.000973 |
| Fibrinogen | 1 | 0.00011 | 5 | 0.002512 | 5 | 0.001638 | 0 | 0 | 5 | 0.001423 |
| Globin/ RBC | 18 | 0.105112 | 19 | 0.100601 | 18 | 0.067832 | 16 | 0.052725 | 21 | 0.109247 |
| Heme/Iron binding | 3 | 0.005258 | 9 | 0.011255 | 7 | 0.011786 | 3 | 0.005854 | 9 | 0.015765 |
| Immunity related | 15 | 0.007953 | 23 | 0.02019 | 17 | 0.014264 | 5 | 0.002246 | 23 | 0.025243 |
| Keratin | 20 | 0.010315 | 34 | 0.012893 | 37 | 0.021782 | 21 | 0.010099 | 40 | 0.045599 |
| Lipocalin | 1 | 0.002122 | 1 | 0.004206 | 1 | 0.00311 | 1 | 0.001514 | 1 | 0.00534 |
| Metabolism, amino acid | 0 | 0 | 1 | 0.000201 | 0 | 0 | 0 | 0 | 1 | 3.83E-05 |
| Metabolism, carbohydrates | 5 | 0.001337 | 9 | 0.001486 | 7 | 0.001096 | 2 | 0.000114 | 7 | 0.001634 |
| Metabolism, energy | 2 | 0.000319 | 2 | 0.000144 | 2 | 0.000364 | 0 | 0 | 6 | 0.000535 |
| Metabolism, lipid | 0 | 0 | 0 | 0 | 0 | 0 | 0 | 0 | 2 | 0.000229 |
| Metabolism, nucleic acids | 1 | 0.000108 | 4 | 0.000989 | 0 | 0 | 0 | 0 | 0 | 0 |
| Nuclear regulation | 19 | 0.013472 | 20 | 0.028935 | 24 | 0.022492 | 10 | 0.008565 | 26 | 0.032846 |
| Protease | 0 | 0 | 4 | 0.000876 | 1 | 9.16E-05 | 0 | 0 | 6 | 0.00066 |
| Protease Inhibitors | 1 | 0.000202 | 3 | 0.000967 | 2 | 0.000151 | 0 | 0 | 7 | 0.00203 |
| Protein export | 3 | 0.001588 | 4 | 0.004882 | 5 | 0.003771 | 2 | 0.00077 | 4 | 0.006886 |
| Protein modification | 6 | 0.000988 | 7 | 0.000754 | 12 | 0.001567 | 0 | 0 | 16 | 0.002522 |
| Protein synthesis | 0 | 0 | 4 | 0.00057 | 4 | 0.00084 | 0 | 0 | 4 | 0.000866 |
| Proteasome machinery | 2 | 0.003209 | 2 | 0.003649 | 2 | 0.003418 | 2 | 0.000584 | 2 | 0.005294 |
| Signal transduction | 8 | 0.003392 | 5 | 0.001237 | 9 | 0.001945 | 0 | 0 | 12 | 0.003857 |
| Transporter/ Receptor | 3 | 0.000381 | 1 | 6.81E-05 | 4 | 0.000521 | 3 | 0.000589 | 13 | 0.00182 |
| Transcription machinery | 0 | 0 | 2 | 6.16E-05 | 2 | 6.68E-05 | 0 | 0 | 7 | 0.000636 |
| Total | 140 | 0.184079 | 199 | 0.230369 | 198 | 0.191521 | 78 | 0.099459 | 282 | 0.311061 |

### The most abundant category of A. americanum tick saliva proteins is tick-specific

Figures 2A and 2B summarizes daily relative abundance of tick saliva proteins during A. americanum tick feeding as determined by normalized spectral abundance factor (NSAF), the index for relative protein abundance (53–55). Fig. 2A shows that three protein categories, tick-specific saliva proteins of unknown function (TSP), protease inhibitors (PI), and heme/iron binding proteins were the most abundant ranging from a respective 24-46%, 11-22%, and 6-26% during feeding (24-192h). Other protein categories at ≤ 10% in abundance include glycine-rich proteins, antimicrobial peptides, evasins, and proteases. For rabbit proteins in A. americanum tick saliva, the most predominant functional category was hemoglobin/red blood cell products (RBC) (13-58%) followed by cytoskeletal (6-20%), heme/iron binding (∼5-16%), keratin (2-30%), and nuclear regulation (2-20%) (Fig. 2B). It is notable that rabbit functional categories related to immunity, antimicrobial peptides, protease inhibitors and proteases were abundant at ≤8% throughout feeding.

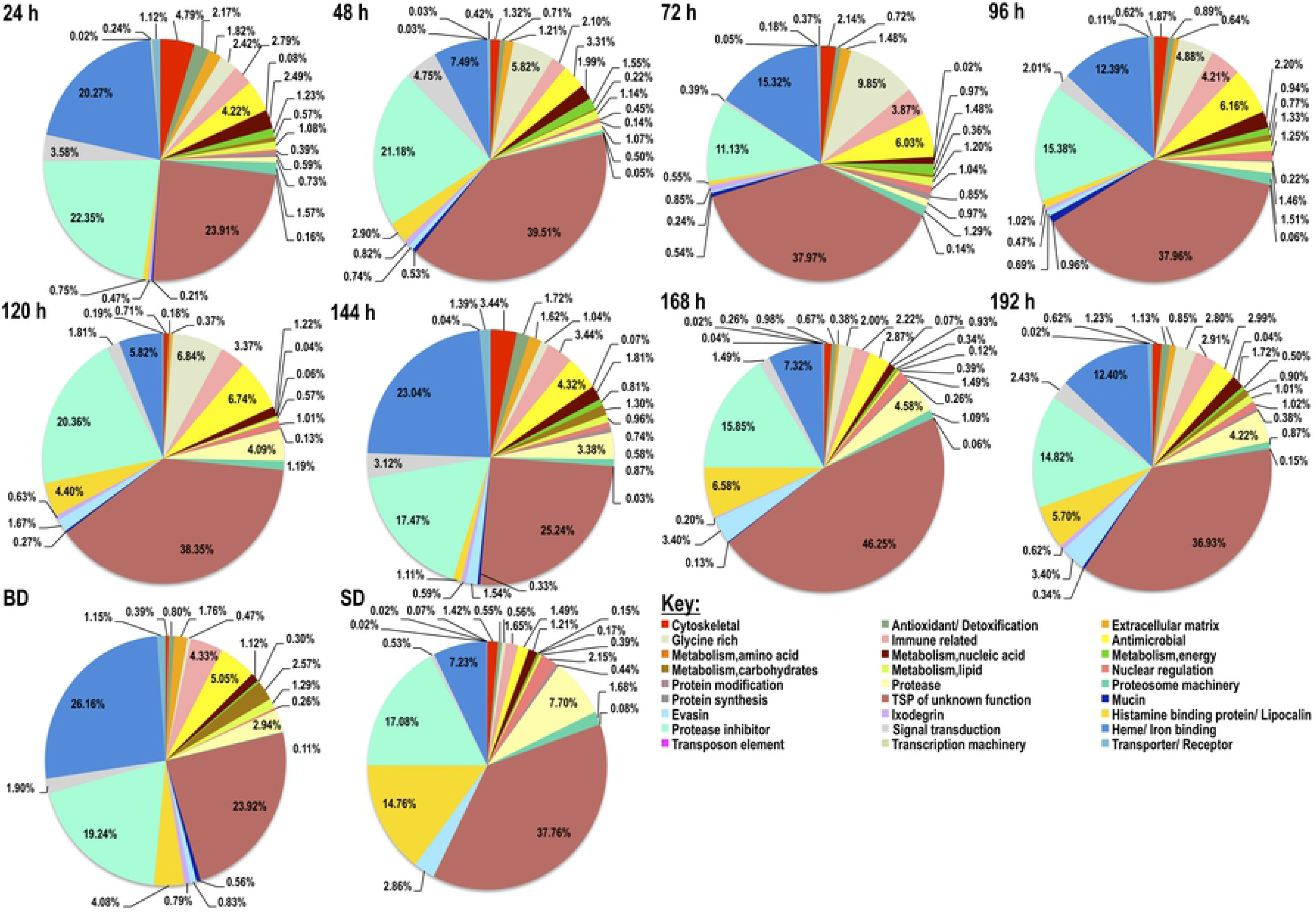
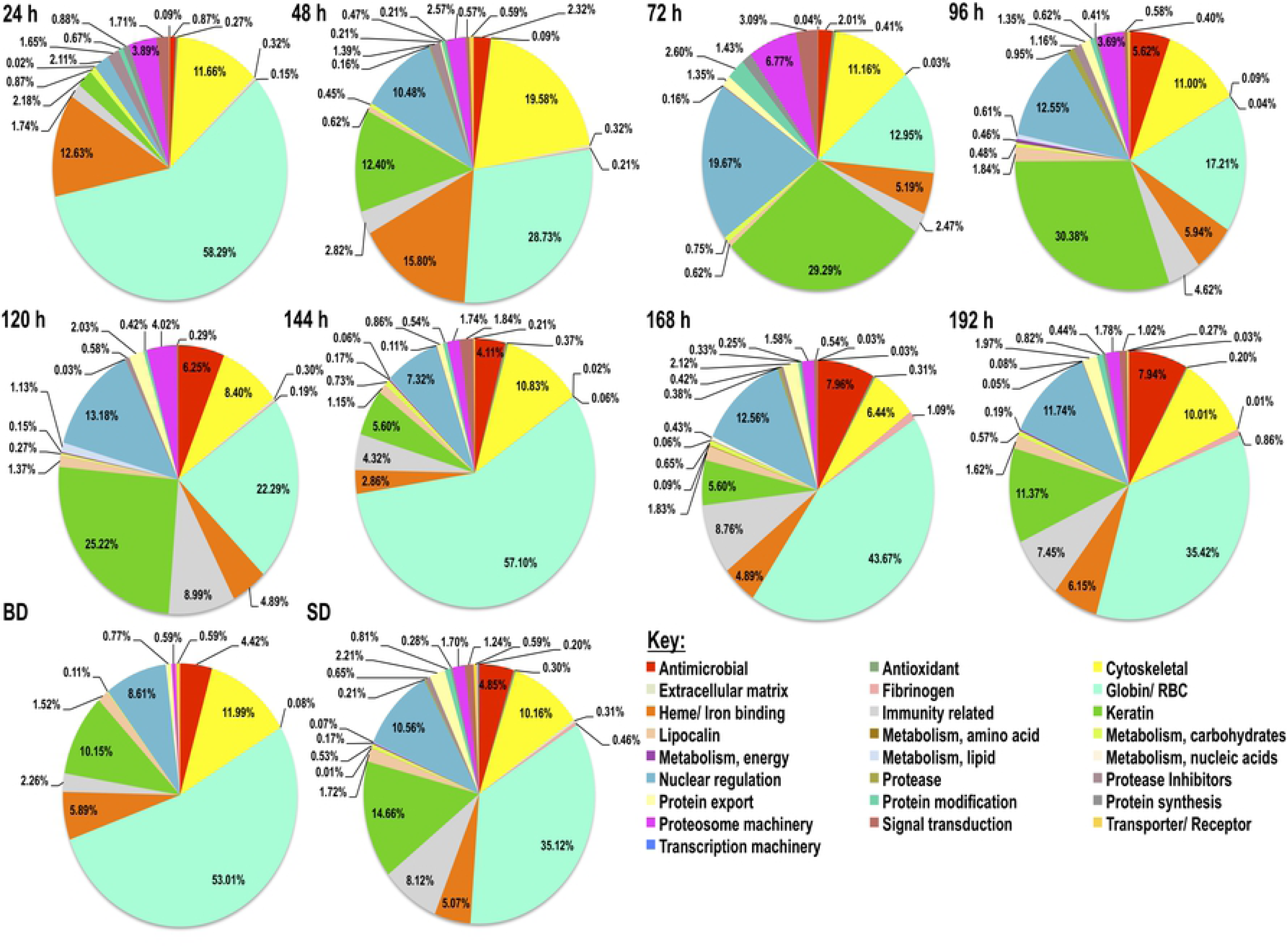

The finding that the majority of proteins identified in this study are of unknown function is not unique to A. americanum tick saliva, it is consistent with findings in saliva of I. scapularis (32) and tick salivary gland transcriptomics (59–61). This is potentially a reflection of how little information exists on the molecular basis of tick feeding physiology.

### Majority of A. americanum tick saliva proteins are associated with early stage tick-feeding processes

To gain insight into broad relationships of secretion dynamics of both tick and rabbit proteins with the tick feeding processes, Z-score statistics normalized NSAF (relative abundance) values were visualized on heat maps (Figs. 3A and 3B). The clustering patterns are influenced by cumulative relative abundance of protein category. The blue to red transition denotes low to high abundance. As shown in figure 3A, the 27 TSP functional categories clustered into four broad secretion patterns (clusters A-D). Broadly, 74% (20/27) of tick protein categories are secreted at high abundance within the first 48 h of feeding (Fig. 3A, clusters A, B, and C) with exception of four categories (evasins, proteases, lipocalins, and nuclear regulation proteins) in cluster D, which are injected into the host at high abundance starting from day five of feeding. These proteins could be important proteins regulating early stages of tick feeding activities such as initiating tick feeding by creating feeding lesion and attaching, while suppressing host tissue repair defenses and also contribute to transmission of TBD agents. Protein categories that were identified in abundance starting from the 192 h feeding time point might be associated regulating the end of the tick-feeding process when the tick detaches from the host skin with minimal damage.

Similarly, the majority of rabbit proteins functional categories (21 of the 25) were detected at high abundance in saliva of A. americanum ticks during feeding (Fig. 3B). The 25 rabbit protein categories in saliva of A. americanum ticks segregated into four clusters, A-D (Fig. 3B). Rabbit proteins that were secreted at high abundance starting from 24-72 h of feeding are part of clusters A and B. Five of the seven proteins in cluster A are highly abundant at 24 and 48 h feeding time points, while those in clusters C and D were less abundant in the first 48 h of feeding and showed varied abundance levels starting from 72 h of feeding.

### Secretion dynamics of non-housekeeping-like A. americanum tick saliva proteins

Supplemental table 2 lists individual proteins that were identified in A. americanum tick saliva. Thirteen functional categories not considered as housekeeping-like (antimicrobial, detoxification extracellular matrix/cell adhesion, evasin, glycine-rich, heme/iron binding, immunity-related, ixodegrin, lipocalin, mucin, protease inhibitors, proteases, and TSPs of unknown function) (Tables 2A and 2B) accounted for 76% of total number of proteins and represented more than 82% in relative abundance throughout feeding time points. In the subsequent sections, we have discussed non-housekeeping-like tick proteins individually per category (Figs. 4A-S) and have highlighted housekeeping-like tick proteins and rabbit proteins as a group below. Our lab is interested and is working to understand functions of proteases and protease inhibitors, and our subsequent discussion below is biased toward these two categories.

#### a) A. americanum tick saliva contains a large diversity of protease inhibitors in nine families

We previously documented at least 18 of the 99 Merops database protease inhibitor (PI) that might be expressed by *A. americanum* and other tick species (62). Here we show that adult *A. americanum* ticks secreted at least 155 PIs belonging into eight PI families. These include Kunitz-type inhibitors (I2, n=68), serine protease inhibitors (serpins, I4, n=21), trypsin inhibitor-like (TIL, I8, n=36), alpha-2-macroglobulins (α2M, I39, n=12), cysteine inhibitors (cystatin, I25, n=12), thyropins (I31, n=3), phosphatidylethanolamine-binding proteins (I51, n=2), and a tick carboxypeptidase inhibitor (TCI, n=1). Of significant interest, nearly 75% of PIs (115/155) in this study were secreted in saliva within the first 120 h post-attached fed ticks (Supplemental table 2). This strongly suggests that functions of tick saliva PIs are associated with regulating early stages of the tick feeding stages such as tick creation of its feeding site and transmission of TBD agents, which are critical to the success of ticks as pests and vectors of TBD agents.

Of the PI families in this study, serpins are the most studied (42, 62–67), presumably because functional roles of this protein category are relatable to tick feeding physiology. To successfully feed and transmit TBD agents, ticks have to overcome serine protease-mediated host defense pathways that are tightly controlled by inhibitors, including serpins. On this basis, it was proposed that ticks might utilize serpins to evade host defenses to successfully feed (68). From this perspective, it is notable that 90% (19/21) of serpins were identified in saliva of ticks that fed for 24-48 h (Fig. 4A), suggesting these serpins are injected into host and might be involved with regulating tick feeding within hours of the tick starting to feed. It is interesting to note that, *A. americanum* serpin 6 and 19, which were previously validated as inhibitors of host defense system proteases (69, 70) were also found in tick saliva within the first 24 h of feeding this study. It is notable that 20-50% of PIs were identified at a single time point for all PI families except serpins, where only 5% (1/21) was found (Supplemental table 2). This might suggest that the functions of tick saliva serpins are important throughout the tick feeding process, most likely in evading host defenses.

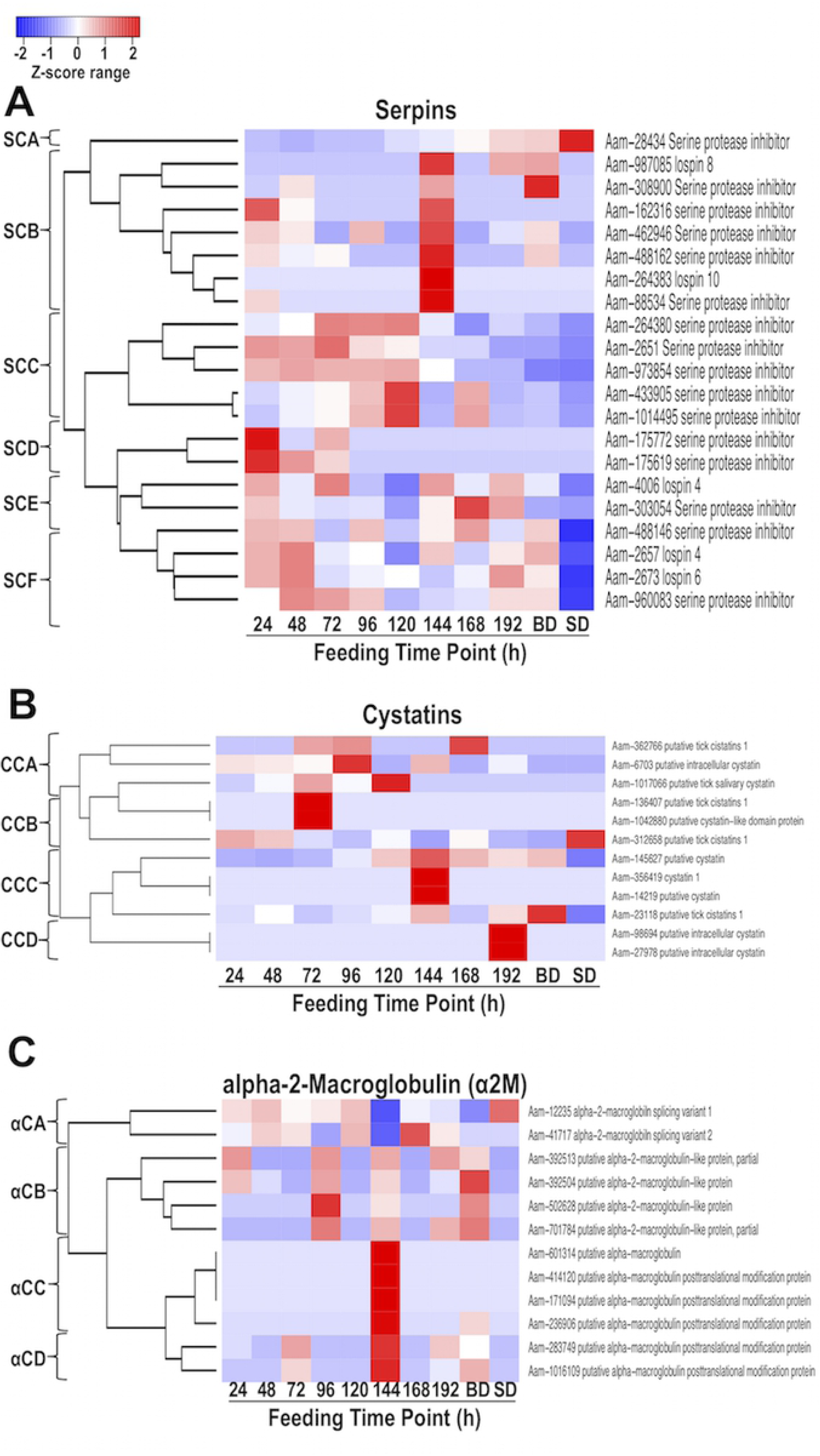

Although not much has been reported on the functional analysis of *A. americanum* tick cystatins, a lone study has reported that RNAi silencing of a cystatin transcript reduced the ability of ticks to feed successfully (71). Several researchers have reported cystatins in other tick species indicated that they play important roles in tick feeding physiology (72). In the soft tick *Ornithodoros moubata,* a cystatin was internalized by host dendritic cells and targeted cathepsin S and cathepsin C, affecting their maturation (73). Cystatins from other tick species also have immunosuppressive functions (74–76). Of the 12 cystatins identified in tick saliva from this study (family I25, Fig. 4B), seven were secreted starting from 72 h of feeding, indicating that majority of cystatins might be involved in regulating tick feeding functions after the tick has initiated feeding. On the contrary, four cystatins were secreted within the first 24 h of feeding and were predominantly secreted throughout feeding.

Similar to cystatins, figure 4C shows the secretion dynamics of alpha-2-macroglobulins (α2M), where majority were injected into the host at high abundance toward the end of feeding (BD). This might suggest that α2M could be involved in regulating tick feeding functions toward the end of tick feeding. There are very few studies on α2M in tick feeding physiology. Two studies have reported the functional roles of α2M in soft tick immune defense (77) and anti-microbial activity in *I. ricinus* (78).

Kunitz-type inhibitors and trypsin inhibitor-like (TIL) were the two most, in number of proteins, identified in all PI families from this study, comprising of a respective 44 and 23% of total PI proteins in tick saliva during feeding. The secretion dynamics of Kunitz-type inhibitors (Fig. 4D) and TIL (Fig. 4E) is comparable and notable: the tick appears to secrete a different set of these inhibitors every 24 h starting from the first day of tick feeding. This might suggest that functions of these inhibitors are required throughout the tick feeding process. It is also notable that a total 35% of Kunitz-type inhibitors and 22% of TILs were detected in saliva of replete fed ticks (SD), unlike the other PIs in this study.

#### b) Majority of proteases in A. americanum saliva are metalloproteases

At the time of this study, protease families that were encoded by *A. americanum* were not enumerated, presumably because its genome had not been sequenced. However, analysis of annotated sequences from *I. scapularis* showed that the tick might encode for all protease categories: aspartic, cysteine, serine, metallo-, and threonine proteases (79). Here, we found that *A. americanum* secretes at least 94 proteases in saliva during feeding. These 94 proteases belong in four categories grouped into 15 families: aspartic (family A1, n=4), cysteine (C1, C2, and C13, n=12), metallo- (M12, M13, M14, M15, M17, M20, M28, and M49, n=56), and serine (S1, S10, and S28, n=22) proteases (Supplemental table 2, Figs. 4F, 4G, and 4H). Please note that the heatmap for aspartic proteases was not developed due to low numbers (the secretion dynamics is presented in Supplemental table 2). The heatmaps (Figs. 4F, 4G, and 4H and Supplemental table 2) show that more than 60% (60/94) of proteases are injected into the host various time points during the first five days of feeding, demonstrating that some of the proteases in this study are associated with tick feeding regulation.

The observation that metalloproteases are the majority of proteases in saliva of *A. americanum* is consistent with our previous findings in the *I. scapularis* proteome (32). It is notable that similar to the *I. scapularis* proteome, metalloproteases that were secreted at high abundance during the first 72 h feeding time points are in families M12 and M13 (Fig. 4G), indicating that these proteases regulate initial tick feeding functions that are important to both tick species. Indirect evidence on snake venom M12 proteases that have anti-coagulant activity (80, 81) suggest that secretion of these proteases at high abundance when the tick is initiating feeding might be beneficial to tick feeding to prevent blood from clotting, which might otherwise prevent blood meal feeding. There is also evidence that RNAi silencing of M12 proteases significantly affected tick-feeding efficiency (82). It has been reported in *I. scapularis* saliva a metalloprotease similar to hemorrhagic proteases of snake venom that act towards gelatin, fibrin(ogen), and fibronectin (83). Likewise, indirect evidence suggests that ticks might utilize M13 proteases to regulate host immunity. In mammals, M13 proteases were among other functions involved in modulating neurotransmitter levels, control blood pressure, involved in reproduction and cancer progression (84).

Another notable similarity between *A. americanum* and *I. scapularis* proteomes is that both tick species secreted a small number of S1 serine proteases, six and three respectively (Fig. 4G and Supplemental table 2). We are interested in S1 serine proteases due to their functional roles in signal transduction as activators of protease-activated receptors (85, 86); could the tick utilize these proteases to interfere with host defense signaling at the tick-feeding site?

The observation that *A. americanum* injected cysteine proteases at the beginning of feeding indicate they might be playing some role(s) in the early stages of tick feeding. Several studies have documented potential functional roles of cysteine proteases in tick physiology (87–89). In a lone study, a cysteine protease from *H. longicornis* when silenced by RNAi, showed to be involved with digestion of a blood meal and increased the number of *Babesia* parasites (90). Recently, a cathepsin L from the tick, *R. microplus* (BmCL1), was shown to interact with thrombin at pH 7.5 and impair thrombin-induced fibrinogen clotting via a fibrinogenolytic activity (91). In helminths, cysteine proteases are the most abundant category of proteins identified into excretion/secretion products (92) and have been shown to be involved with host immune evasion (93) and extracellular matrix degradation (94).

Majority of studies on tick aspartic proteases are mainly characterized as blood digestion proteins in the midgut, similar to the mammalian lysosome acidic protease, cathepsin D (95). In *H. longicornis* adult ticks, the potential role of these proteins in proteolysis of erythrocyte hemoglobin has been reported (96). Other studies have shown the importance of this protease in embryogenesis, playing roles in vitellin degradation (97) and heme-binding properties (98). Although only four aspartic proteases were identified in *A. americanum* saliva during feeding, three of these proteases were present within the first 96 h of feeding, which may implicate roles in the early stages of tick feeding success (Supplemental table 2).

#### c) Lipocalins/histamine-binding proteins are alternately secreted during tick feeding

Inflammation response is among host defense pathways that ticks must evade to complete feeding. Histamine is one of the key mediators of inflammation in tissue damage that is expected to occur in response to tick feeding (99). From this perspective, lipocalins/tick histamine-binding proteins in tick saliva are suspected to be part of the tick machinery to evade the host’s inflammation defense response through sequestration of histamine that is released at the tick-feeding site. In this study, we found 46 lipocalins/tick histamine-binding proteins that show two broad secretion patterns: secreted at multiple feeding time points and those that were alternately secreted at single time points (Fig. 4I). It is interesting to note that of the total 46 lipocalins identified in tick saliva during feeding, 22% (10/46) were present within the first 48 h of feeding, while 35% (16/46) were present after 96 h of feeding, and 43% (20/46) were identified in a single time point (Fig 4I, Supplemental table 2). Given that in addition to regulating inflammation, lipocalins/histamine-binding proteins have other diverse functions such as antimicrobials (100, 101), glucose metabolism (102) and binding several ligands including serotonin and fatty acids (103, 104), it is most likely that these proteins might be involved in regulating several other tick feeding functions besides mediating the tick’s anti-inflammation function.

#### d) Heme binding proteins are secreted at high abundance throughout feeding

Like other animals, ticks require iron and heme (the iron-containing part of hemoglobin) for normal physiological functions (105). However, ticks do not have a heme biosynthesis pathway, therefore they must obtain it from host blood (106). Female ticks that were artificially fed a diet not containing hemoglobin laid sterile eggs (107) demonstrating the importance of heme in tick biology. However, in high abundance heme can be toxic for the tick (108), therefore it is postulated that hemelipoproteins and vitellogenenins could serve as heme binding proteins to remove the excess heme from the tick system. Supplemental table 2 lists a total of 17 heme/iron-binding proteins consisted of hemelipoproteins, vitellogenins, and a ferritin that collectively accounted for the third most abundant protein category in tick saliva throughout feeding (Fig. 2). High abundance of hemelipoproteins here is in consistent with other tick saliva proteomes (32–34). The secretion dynamics summarized in figure 4J revealed two broad secretion patterns, those that are injected into the host from 24 h through 120 h of feeding (HCB and HCC) and those that injected into the host starting from 144 h of feeding through the end of tick feeding (HCD and HCA). Ticks acquire both iron and heme from host blood (106, 109), and thus iron and/heme-binding proteins are important to normal tick physiology. It has been shown that *R. microplus* hemelipoprotein (HeLp) could bind eight heme molecules (110). Given that hemelipoprotein is the most abundant protein in tick hemolymph (111), it could be secreted in saliva as a result of this protein being transferred into the salivary glands when exposed to the hemolymph. However, transcriptional profile and protein localization of these hemelipoproteins in salivary glands of unfed and fed adult ticks suggest that they could act in different pathways during blood-feeding (112). Among other functions it is known that free heme has pro-inflammatory properties (113). Thus, the presence of hemelipoproteins could lower free heme concentration at the feeding site, reducing inflammation. Other roles of tick hemelipoproteins such as an antioxidant in transporting other compounds such as cholesterol, phospholipids, and free fatty acids have been previously reported (114). It is interesting to note that reduction of vitellogenin receptor (VgR) expression by RNAi resulted in reduced fertility (115) and *Babesia bovis* transmission and oocyte maturation (116).

#### e) Ticks inject multiple antioxidant proteins into the feeding site

Feeding and digestion of large amounts of host blood exposes ticks to hydroxyl radicals and reactive oxygen species (ROS), which if left uncontrolled could damage tick tissue (117, 118). Expression of antioxidant proteins protect the tick during feeding and digestion of the blood meal. Studies have shown that RNAi silencing of tick antioxidants caused deleterious effects to the tick and prevented them to obtain a full blood meal (119, 120). Previous studies by others and from our lab have documented presence of antioxidants in tick salivary glands (121, 122) and saliva (32-34, 123, 124). In this study we identified 41 putative antioxidant enzymes. These enzymes include glutathione-S-transferase, thioredoxin, superoxide dismutase, catalase, peroxinectin, arylsulfatase, aldehyde dehydrogenase, epoxide hydrolase, sulfotransferase, sulfhydryl oxidase and glycolate oxidase. Figure 4K reveals two broad secretion patterns of tick saliva antioxidant proteins based on NSAF values as an index for abundance: (i) proteins injected into the host in high abundance once at various feeding time points (ACA-ACI) and (ii) proteins that are consecutively injected into the host in high abundance from 24-96 h of feeding (ACF). Like heme/iron binding proteins, tick antioxidants are presumed to function inside the tick; the question is why do ticks inject these into the feeding site? Host tissue injury caused during the creation of the tick-feeding site could trigger release of oxidants such as ROS; could tick saliva antioxidants function to cleanse the blood meal before the tick ingests it or potentially to protect the host from damage to keep the feeding site balanced?

#### f) Glycine-rich and extracellular matrix/cell adhesion proteins are secreted early during tick feeding

Within 5-30 min of attachment, the tick secretes an adhesive substance called cement, which anchors ticks onto host skin during its protracted feeding period (125). Tick cement is also suggested to protect the tick from host immune factors (126, 127) and might function as antimicrobials at the feeding site (128). Glycine-rich proteins are among categories of tick proteins that are thought to play key roles in formation of tick cement (125). From this perspective, glycine-rich proteins are among tick proteins that have received significant research attention (129–132). In this study we found a total of 67 glycine-rich proteins, which represented the fifth largest category of proteins identified in tick saliva during feeding (Fig. 2). Nearly 90% (60/67) of the glycine-rich proteins were secreted in abundance within the first four days of feeding (Fig. 4L; GCB, GCD, GCE, GCF, and GCG). Tick cement deposition is completed during the first 96 h of tick feeding (125), and thus it is conceivable that some of the glycine-rich proteins in this study might be involved with tick cement formation. It is interesting to note that some of the glycine-rich proteins that were identified from tick cement in our lab (131) and others (130) were also found in this study (Supplemental table 3). Some of the glycine-rich proteins were secreted from the 144-h time point, long after tick cement formation; these might regulate other tick feeding functions. Although glycine-rich proteins are mostly known for their potential role in tick cement formation, indirect evidence in other organisms indicate that these proteins might be involved in other functions such as host defense and stress response as in plants (133).

Figure 4M summarize the secretion dynamics of 37 extracellular matrix proteins that were found in this study. Similar to glycine-rich proteins, majority (27/37) of the extracellular proteins were secreted within the first five days of feeding demonstrating their role in early stage tick feeding regulation. Our speculation is that some of these proteins will play roles in formation of tick cement. In a previous study, RNAi silencing of chitinase, also identified in this study, weakened the tick cement cone to the extent that host blood was leaking out around the mouthparts of attached ticks (40).

#### g) Antimicrobials, mucins, and immune related proteins are secreted throughout the feeding process

Once the tick has anchored itself onto the host skin and created its feeding lesion, it faces a difficult task of overcoming host humoral and cellular immunity, and also preventing microbes in the host skin from colonizing the tick-feeding site. Here we show that *A. americanum* secretes immunomodulatory and antimicrobial peptides starting within the early stages of the tick feeding process (Fig. 4N-R). We identified nine antimicrobials consisting of microplusins, lysozymes, and defensins (Fig. 4N). Previous studies showed that microplusin has dual effects against fungus and gram-positive bacteria, lysozyme against gram-positive bacteria, and defensin effective against both gram-positive and -negative bacteria (134–136). The heat map in figure 4N shows that antimicrobials were injected into the host starting at 24 and 48 h (AMCA), from 72 h (AMCC), and from 120 h (AMCB). This secretion pattern suggests that the functions of antimicrobial peptides are needed throughout feeding.

Similar to antimicrobials, we identified 12 mucins (Fig. 4O), with ∼60% of these proteins (7/12) being secreted at high abundance within 24-48 h of feeding. Functional roles of mucins in ticks have not been studied. However, indirect evidence in mammals suggest that mucins might be involved in antimicrobial activity in that human mucins were shown to encapsulate microbes (137).

Among putative immunomodulatory proteins, we identified evasins (Fig. 4P) and ixodegrins (Fig. 4Q). Evasins (n=12, Fig. 4P) were shown to bind to chemokines (138, 139) to reduce leukocytes recruitment to the tick feeding site and therefore contribute to tick evasion of the host’s inflammatory defense. It is interesting to note that the 12 evasins identified in tick saliva were present after 24 h of feeding and continued to be secreted throughout feeding at variable levels. This might suggest that evasins might not be involved in regulating tick feeding functions during the first 24 h of tick feeding.

Figure 4Q summarizes the secretion pattern of the six ixodegrin-like proteins found in tick saliva during feeding in this study. It is interesting to note, 83% (5/6) of these proteins were identified within the first 48 h of feeding (Supplemental table 2.) These proteins were first described in *I. scapularis* as inhibitors of platelet aggregation (140). Platelet aggregation is the first step in the blood clotting system (141), which ticks must overcome to successfully feed. Thus, the presence of ixodegrins in saliva of *A. americanum* at the start of feeding is beneficial to tick feeding success. Finally, we also found proteins that show similarity to previously characterized immunomodulatory proteins (Fig. 4R), which have been validated in other tick species including p36, which inhibits cell proliferation and cytokine expression (142). These proteins might play roles in mediating the tick’s evasion of host immunity.

#### h) Tick-specific secreted saliva proteins (TSP) of unknown function are alternately secreted

Over one-third of Ixodidae protein sequences deposited into GenBank are annotated as hypothetical, secreted, conserved and unknown proteins. However, some are annotated based on sequence identities and conserved signature motifs, which include basic tail/tailless proteins, 8.9 kDa protein family, leucine-rich proteins, AV422 (a tick saliva protein that is high upregulated when ticks are stimulated to start feeding [39, 143]), proteins containing RGD motifs, which might play roles in inhibition of platelet aggregation (140, 144). In this study we have identified a total of 377 (Fig. 4S) tick saliva proteins that fit the above description that we refer here to as tick-specific saliva proteins of unknown functions (TSPs). More than 95% (357/377) of the total TSPs were identified within the first eight days of feeding in tick saliva indicating their potential roles in regulating the tick feeding process. It is interesting to note that the secretion pattern for over a third (128/377) of the total TSPs identified in tick saliva during feeding were alternately injected once during feeding (Supplemental table 2). From the perspective of finding target antigens for tick vaccine development, TSPs represent a unique opportunity in that they do not share any homology to host proteins and might not cross-react with the host.

### A. americanum secretes multiple housekeeping-like proteins in saliva throughout the feeding process

Supplemental table 2 lists 288 housekeeping-like proteins that were identified in this study. Presence of these proteins in *A. americanum* saliva is not unexpected, as similar findings have been previously reported in tick saliva (32–34). The 288 housekeeping-like proteins were classified into 14 categories including those associated with metabolism of amino acids (n=7), carbohydrates (n=25), energy (n=31), lipids (n=31), and nucleic acids (n=33). Other protein categories include those involved in cytoskeletal (n=53), nuclear regulation (n=16), protein modification (n=21), proteasome machinery (n= 8), protein synthesis (n=10), signal transduction (n=24), transposable element (n=3), transcription machinery (n=7), and transporter/receptors (n=17). It is interesting to note that, within the first 24 h of feeding 12 of the 14 categories were identified at high abundance (Fig. 3A).

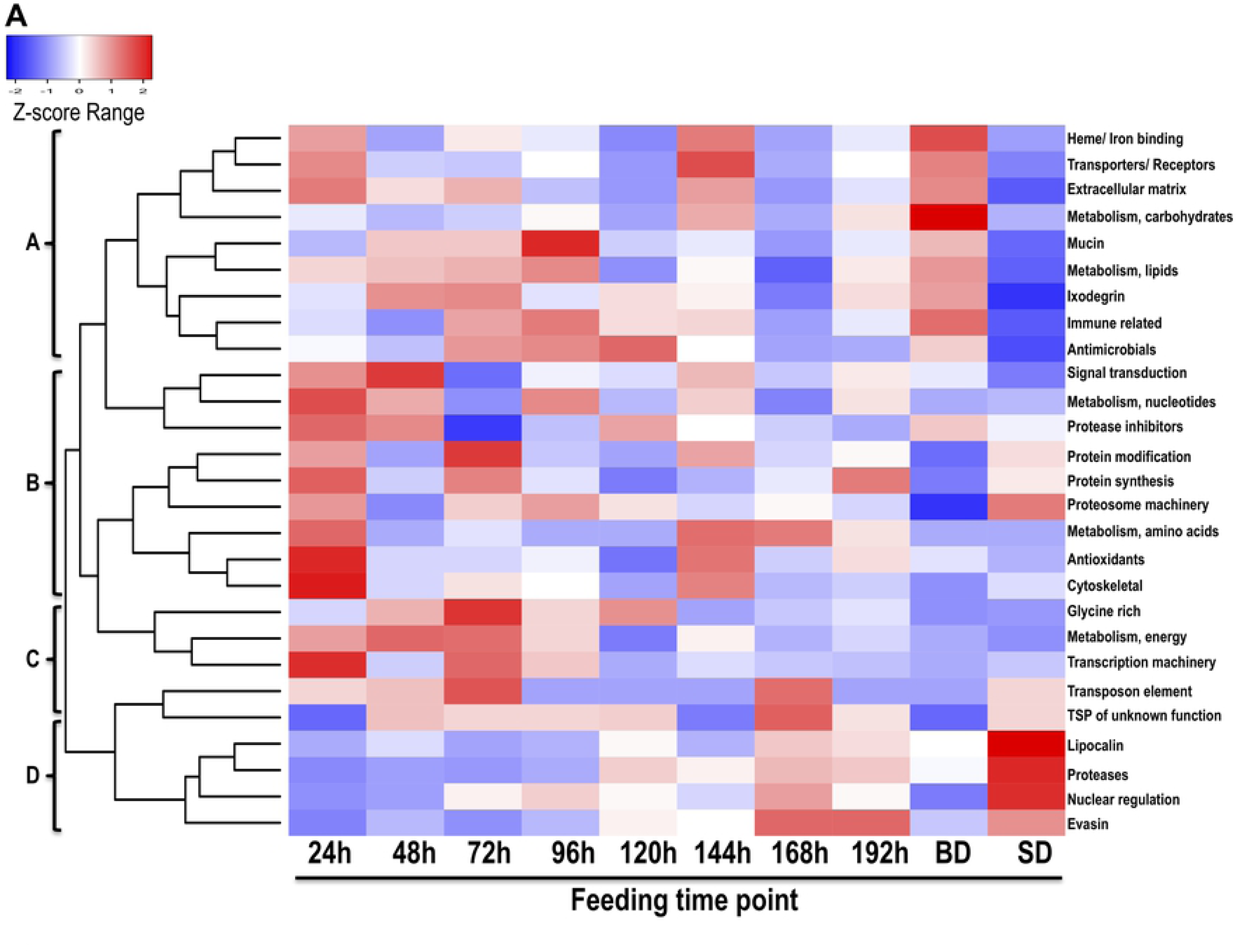
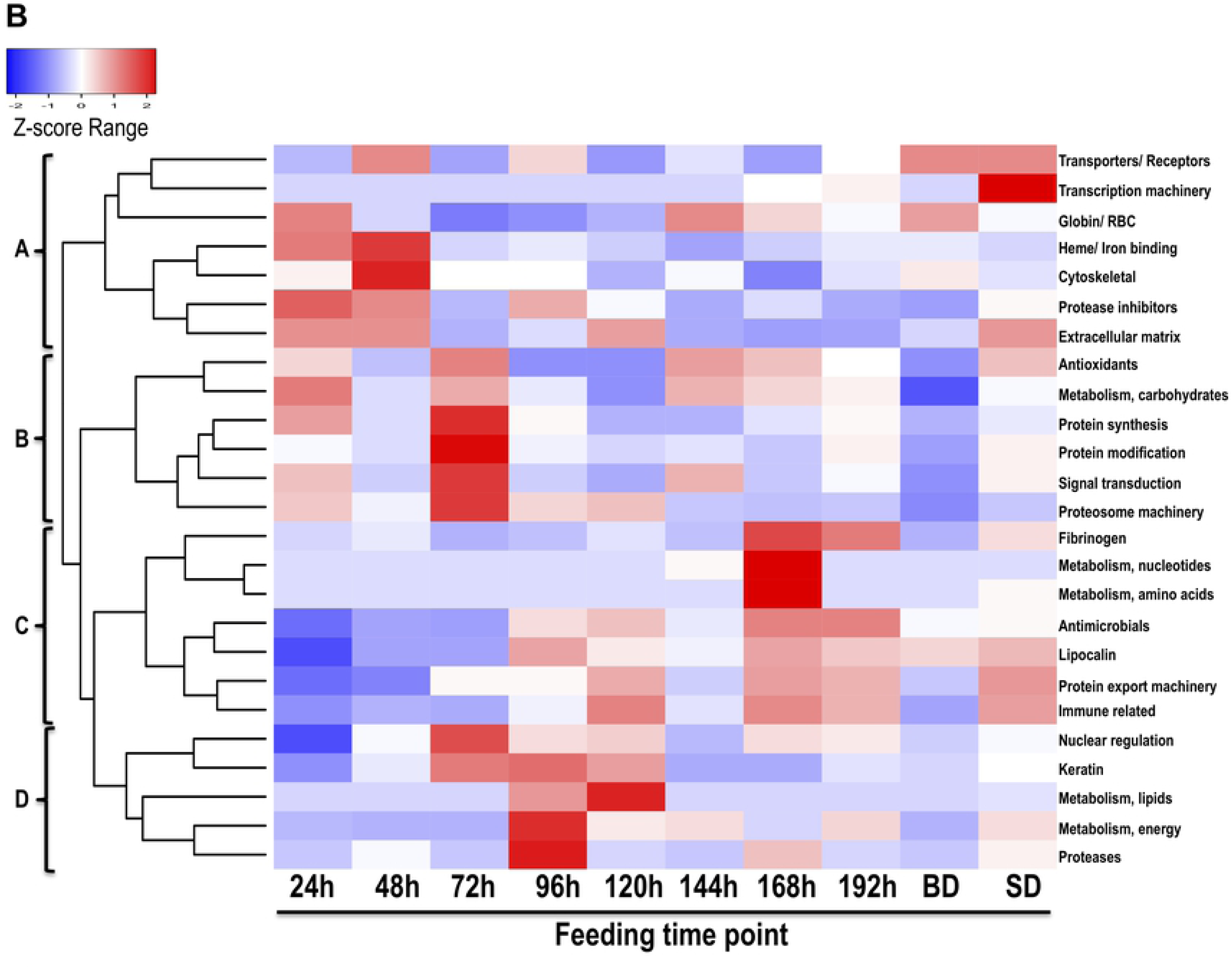

One feature of housekeeping-like tick proteins is that they have high sequence identity with mammalian housekeeping proteins, and for this reason they are discounted as potential target antigens for tick vaccine development. However, based on literature showing that several roles of these proteins in host defense, we think that these proteins play an important role in tick feeding physiology. Housekeeping-like proteins identified here mostly function intracellularly, and they serve as alarm signals to alert the host defense system to injury when secreted outside of the cell (145). There is evidence that in the extracellular space, some of the housekeeping proteins such as heat shock proteins, have anti-inflammatory functions (146), while histone proteins have antimicrobial activity (147). Given high sequence similarity to host housekeeping proteins, could it be that some of the tick housekeeping-like proteins play roles in promoting tick feeding through anti-inflammatory and anti-microbial activity?

Another important aspect of tick feeding physiology that has not received much attention is the fact that host blood meal also contains carbohydrates, lipids and other molecules besides host proteins. It is notable that some of the tick housekeeping-like proteins in tick saliva have high similarity to enzymes in the carbohydrate and lipid metabolism pathways. Is the tick pumping these proteins into the feeding site to process these molecules before the tick takes its blood meal?

### Secretion of rabbit host proteins in A. americanum tick saliva is not random

In this study, we identified 335 rabbit host proteins belonging into 25 different categories that include cytoskeletal (19%), keratin (13%), nuclear regulation (8%), immune-related (8%), hemoglobin/RBC degradation (6%), transporters/receptors (5%), protein modification (5%), and protein categories below 4% included antimicrobials, extracellular matrix, heme/iron binding, detoxification/ antioxidants, metabolism (energy, carbohydrates, lipid, amino acid, and nucleic acids), protein export, protein synthesis, fibrinogen, protease inhibitors, proteases, signal transduction, transcription machinery, proteasome machinery, and lipocalin (Tables 3A and 3B, Supplemental table 2). Relative abundance as determined by NSAF indicated that the most abundant protein categories consisted of hemoglobin/RBC degradation products (58-13%), followed by heme/ iron binding host proteins (13-16%), and cytoskeletal (6-20%) (Fig. 3).

At a glance, presence of rabbit host proteins in *A. americanum* tick saliva could be dismissed as host protein contamination. This observation might be strengthened by the fact that some rabbit host proteins in tick saliva such as keratin, nuclear regulation proteins, and host antimicrobial peptides increased in abundance as feeding progressed. This suggested that secretion of host proteins into tick saliva was a consequence of ticks ingesting an increased amount of host blood, and that some of these host proteins might leak or be regurgitated back into the host via saliva or esophagus. However, our data here suggests that the tick might systematically be utilizing host proteins to regulate its tick-feeding site. For instance, mammals are likely to encode for more than 500 proteases and 150 protease inhibitors (based on rat, mice, and humans [148]), however we found 9 proteases and 8 protease inhibitors from host origin in *A. americanum* tick saliva (Supplemental table 2). We are of the view that if secretion of host proteins was a random process, we could have identified more rabbit host proteases and protease inhibitors. There are reports that human α1-antitrypsin and α2-macroglobulin are secreted following injury as occurs during tick feeding, and if left uncontrolled could lead to delayed wound healing (149), which is beneficial to tick feeding. On this basis, it is highly likely that ticks inject host α1-antitrypsin and α2-macroglobulin into the feeding site as a strategy of evading the host’s tissue repair defense response. It is also notable that fibrinogen and neutrophil gelantinase-associated lipocalin, which among other functions plays important roles in wound healing, were identified towards the end of feeding (150–153). This is interesting in that the tick-feeding lesion is completely sealed, preventing leakage of blood, when a replete fed tick detaches from its feeding site. It has been reported in *Opisthorchis viverrini*, the human liver fluke, that they secrete proteins in the granulin family that accelerate wound healing (154). Could it be that the increased abundance of host proteins involved in wound healing are secreted by the tick into the feeding site towards the end of tick feeding is the tick’s way to help its host heal?

### Different tick species might utilize similar proteins to regulate feeding

At the time of preparing data in this study for publication, several other tick saliva proteomes had been published. We took advantage of the availability of these data to test the hypothesis that key proteins that are important to tick feeding might be conserved across tick taxa. Thus, we compared data in this study to saliva proteomes of *I. scapularis* (32), *R. microplus* (33), *H. longicornis* (34), *R. sanguineus* (35), *D. andersoni* (36), and *O. moubata* (37). This analysis revealed that more than 24% (284/1182) of the *A. americanum* tick saliva proteins in this study have homologs in saliva proteomes of other tick species (Supplemental table 4). Table 4 highlights the 163, 138, 137, 92, 22, and 11 *A. americanum* tick saliva proteins in 22 categories that were >70% identical to proteins in saliva of *I. scapularis* (32), *H. longicornis* (34), *D. andersoni* (36), *R. microplus* (33), *O. moubata* (37) and *R. sanguineus* (35), respectively. Of the 22 categories of proteins, immune-related proteins were present in all tick saliva proteomes. Likewise, proteins from nine other categories (antioxidant/detoxification, carbohydrate metabolism, cytoskeletal, extracellular matrix, heme/iron binding protease, protease inhibitor, protein modification, and signal transduction) from *A. americanum* saliva were present in five other tick saliva proteomes. It is interesting to note when comparing saliva proteins in the *A. americanum* tick-specific saliva secreted protein category with other hard ticks, with the exception of *R. sanguineus*, five proteins were shared with Prostriata *I. scapularis*, and 6, 22, and 14 from Metastriata *H. longicornis*, *D. andersoni*, and *R. microplus*, respectively.

**Table 4.** *Amblyomma americanum* tick saliva protein categories that are conserved in other tick saliva proteomes at 70% identity

| Classification | *I. scapularis* | *H. longicornis* | *D. andersoni* | *R. microplus* | *O. moubata* | *R. sanguineus* |
| --- | --- | --- | --- | --- | --- | --- |
| Cytoskeletal | 36 | 27 | 19 | 12 | 3 | 0 |
| Detoxification | 13 | 9 | 9 | 5 | 0 | 2 |
| Extracellular matrix | 3 | 6 | 9 | 3 | 0 | 1 |
| Glycine rich | 5 | 4 | 4 | 4 | 0 | 0 |
| Immune related | 4 | 3 | 3 | 4 | 1 | 1 |
| Metabolism, amino acids | 4 | 1 | 0 | 0 | 0 | 0 |
| Metabolism, carbohydrates | 4 | 3 | 6 | 1 | 1 | 0 |
| Metabolism, energy | 20 | 11 | 12 | 0 | 4 | 0 |
| Metabolism, lipids | 2 | 1 | 3 | 4 | 0 | 0 |
| Metabolism, nucleic acids | 11 | 9 | 1 | 0 | 2 | 0 |
| Nuclear regulation | 6 | 5 | 6 | 4 | 0 | 0 |
| Protein modification | 16 | 14 | 8 | 9 | 7 | 0 |
| Protease | 6 | 6 | 8 | 3 | 0 | 0 |
| Proteasome machinery | 7 | 6 | 5 | 0 | 0 | 6 |
| Protein synthesis | 4 | 2 | 3 | 2 | 0 | 0 |
| Secreted saliva proteins | 5 | 6 | 22 | 14 | 0 | 0 |
| Lipocalin | 0 | 0 | 0 | 4 | 0 | 0 |
| Protease Inhibitors | 5 | 14 | 10 | 11 | 1 | 1 |
| Signal transduction | 6 | 3 | 6 | 1 | 1 | 0 |
| Heme/iron binding | 1 | 7 | 2 | 9 | 2 | 0 |
| Transcription machinery | 2 | 0 | 1 | 1 | 0 | 0 |
| Transporters/ receptors | 3 | 1 | 0 | 1 | 0 | 0 |
| Total Protein Matches | 163 | 138 | 137 | 92 | 22 | 11 |

We would like the reader to note that with the exception of *I. scapularis* tick saliva proteome for which proteins were identified every 24 h during feeding (32) the other tick saliva proteomes were limited to a narrow range of tick feeding time points and/or fully engorged ticks. This might be the reason that higher numbers of *A. americanum* tick saliva proteins were found compared to *I. scapularis* tick saliva proteome. It is interesting to note that, *A. americanum* and *I. scapularis* are biologically different as they belong to different tick lineages, Prostriata and Metastriata (155, 156). Thus, tick saliva proteins that are shared between these two tick species could regulate evolutionarily conserved proteins that regulate essential tick feeding physiology functions. On this basis, such proteins could be targeted for tick vaccine development. We have previously shown that RNAi silencing of *A. americanum* tick saliva serpin 19, an anti-coagulant (69), which is also conserved in *I. scapularis* ticks (42, 62), caused significant mortality demonstrating the importance of this protein in tick physiology (32).

## Conclusion and Future perspective

This study has made a unique contribution toward understanding the molecular basis of *A. americanum* tick feeding physiology. We believe that this study provides a good starting point toward discovery of effective targets for anti-tick vaccine development. Our strategy to identify tick saliva proteins every 24 h during feeding has allowed us to map tick saliva proteins to different phases of the tick feeding process. This is significant as it provides for the opportunity to focus on tick saliva proteins that regulate the tick feeding process that precede critical events such as TBD agent transmission. Majority of TBD agents are transmitted after 48 h of tick attachment (157, 158), and therefore proteins that are secreted from 24 and 48 h of tick feeding time points are prime candidates for tick vaccine research. It is important to acknowledge the fact that, during the course of feeding, *A. americanum* ticks secretion of more than 1500 tick and rabbit host proteins might indicate that the tick has inbuilt systems to evade host immunity, and that it is going to be a challenge to actually find effective targets for anti-tick vaccine development. However, the findings that nearly 300 *A. americanum* tick saliva proteins were also secreted by other tick species is very encouraging as these proteins might provide insight into conserved mechanisms that are utilized by all ticks to successfully feed, and could serve as potential targets for anti-tick vaccine development.

We have recently described proteins (n=340) in saliva of unfed *A. americanum* ticks that were stimulated to start feeding on three different hosts: rabbits, dogs, and humans (38). It is notable that 70% (231/340) of proteins in saliva of unfed *A. americanum* ticks were found in the tick saliva proteome described here (Supplemental table 5). The significance of these data is that the 231 tick saliva proteins present in saliva of both unfed and fed ticks represent proteins that are potentially injected into the host within minutes of the tick attaching onto host skin and are likely associated with regulating initial tick feeding events. Immunologically blocking functions of these proteins might significantly disrupt tick feeding and prevent transmission of TBD agents. In summary, this study has set the foundation for in-depth studies to understand *A. americanum* tick feeding physiology and find effective targets for development of tick-antigen based vaccines to prevent TBD infections.

## Acknowledgements

This research was supported by National Institutes of Health grants (AI081093, AI093858, AI074789, AI074789-01A1S1) to AM and National Center for Research Resources (5P41RR011823) and National Institute of General Medical Sciences (8P41GM103533) to JRY.

The funders had no role in study design, data collection and analysis, decision to publish, or preparation of the manuscript.

## Data Availability

The mass spectrometry proteomics data have been deposited to the ProteomeXchange Consortium via the PRIDE (159) partner repository with the dataset identifier PXD014844.

